# Multisensory integration of social signals by a pathway from the basal amygdala to the auditory cortex in maternal mice

**DOI:** 10.1101/2022.02.17.480854

**Authors:** Alexandra C. Nowlan, Jane Choe, Hoda J. Ansari, Clancy Kelahan, Stephen D. Shea

**Affiliations:** Cold Spring Harbor Laboratory, 1 Bungtown Road Cold Spring Harbor, New York 11724, USA

## Abstract

Social encounters are inherently multimodal events, yet how and where social cues of distinct sensory modalities merge and interact in the brain is poorly understood. When their pups wander away from the nest, mother mice use a combination of vocal and olfactory signals emitted by the pups to locate and retrieve them. Previous work revealed the emergence of multisensory interactions in the auditory cortex (AC) of both dams and virgins who co-habitate with pups (‘surrogates’). Here we identify a neural pathway that integrates information about odors with responses to sound. We found that a scattered population of glutamatergic neurons in the basal amygdala (BA) projects to the AC and responds to odors, including the smell of pups. These neurons exhibit increased activity when the female is searching for pups that terminates upon contact. Finally, we show that selective optogenetic activation of BA-AC neurons modulates responses to pup calls, and that this modulation switches from predominantly suppressive to predominantly excitatory after maternal experience. This supports an underappreciated role for the amygdala in directly shaping sensory representations in an experience-dependent manner. We propose that the BA-AC pathway integrates olfaction and audition to facilitate maternal care, and speculate that it may carry valence information to the AC.

## INTRODUCTION

In sharp contrast to how most sensory neurophysiology and psychophysical research has historically been conducted, organisms engaged in unstructured natural behavior typically make decisions by considering all sensory data available to them. For example, during social encounters, individuals are often informed about the identity and status of conspecifics by interpreting a combination of auditory, olfactory, tactile, and/or visual cues. While the significance of all these modalities have individually been well-studied, much less is known about how qualitatively distinct pieces of social information from different senses are integrated in the brain to guide behavior. Here we examine the neural circuitry that underlies multisensory integration of odor and sound during maternal behavior in mice.

Shortly after giving birth, first time mother mice learn to respond to ultrasonic vocalizations (USVs or ‘calls’), emitted by pups when they are separated from the litter, by locating them and bringing them back to the nest (‘pup retrieval’) [1, 2]. Sensory experience with pups is sufficient to motivate pup retrieval because virgin females co-housed with pups (‘surrogates’) also learn to retrieve without the influence of pregnancy hormones [3–10]. Consistent with the importance of USV detection for pup retrieval, activity and plasticity of the auditory cortex (AC) is implicated in accurate and efficient pup retrieval [5–9]. Audition and olfaction are jointly required because disrupting olfactory processing also interferes with retrieval [8, 11–13]. Interestingly, there is evidence that separate signals related to odor and sound may merge in the AC. Delivery of pup odors to anesthetized female mice acutely modulates single neuron AC responses to a range of sounds including USVs, but only in pup-experienced females (mothers and surrogates), not naïve females [8, 9]. The brain pathway by which odor information reaches the AC is unknown.

We took advantage of the ability of virgin females to learn pup retrieval in order to study the neural circuit that integrates odor signals in the AC. This allowed us to observe the effects of sensory experience on this pathway independent of pregnancy hormones. We report the following findings. First, we used anatomical tracing to identify a population of neurons in the basal amygdala (BA) that project into the AC. Second, we used an intersectional viral strategy to label the AC projecting BA neurons (BA-AC) with the calcium (Ca^2+^) sensor GCaMP6s. Fiber photometry in awake, head-fixed mice revealed that BA-AC neurons respond to odor, including the smell of pups. Third, we performed fiber photometry in freely behaving mice and found that their activity is elevated during active search for pups. Finally, we optogenetically activated BA-AC neurons in anesthetized females and found that activation modulates responses to sound. This modulation switches from predominantly suppressive to predominantly excitatory after maternal experience. Based on these observations, we propose that neurons in the BA carry odor information to the AC, where they influence auditory activity to improve pup retrieval. This contrasts with other forms of multisensory integration in the auditory cortex that emerge via inputs from primary sensory structures. Moreover, our results reveal an underappreciated role for the amygdala in directly modulating sensory representations in accordance with maternal status.

## RESULTS

### Olfaction is required for pup retrieval behavior in surrogate females

Nulliparous female mice that have sustained exposure to pups, e.g. via co-habitation with a mother and her litter, learn to perform efficient pup retrieval within a few days [3, 4]. Previous reports regarding the influence of pup odors on auditory activity in the AC found evidence of this integration in both mothers and surrogates. Here we took advantage of this property by performing all experiments reported here in surrogates to focus on the role of sensory experience in shaping auditory flexibility and maternal behavior.

The necessity of olfactory signals for maternal behavior in dams has already been established [8, 11–13], however it is unknown whether surrogates share this requirement. Thus, we performed behavioral experiments to test the importance of odor cues for retrieval in surrogates. We assayed the pup retrieval performance of naïve virgin females beginning prior to co-habitation with a pregnant dam and continuing throughout their surrogacy on postnatal days zero, three, and five (P0, P3, P5, Figure 1A, B). Performance was quantified using a normalized measure of latency to retrieve (‘latency index’; see Materials and Methods) [6]. Then, we used the tissue specific toxicant methimazole (MMZ) to ablate the olfactory epithelium of these mice and measured their retrieval performance again (Figure 1F). As expected, naïve virgins exhibited poor pup retrieval (naïve latency index: 0.622 ± 0.09; all values are reported as mean ± SEM unless otherwise noted). The same mice were each subsequently paired with a pregnant dam as surrogates, and they exhibited a significant decrease in retrieval latency on P3 (P3 latency index: 0.060 ± 0.01; Tukey’s multiple comparisons test; *p* = 0.023) and P5 (P5 latency index: 0.052 ± 0.01; Tukey’s multiple comparisons test; *p* = 0.017) compared to their naïve performance (Figure 1C). After MMZ treatment, retrieval latency substantially increased (MMZ latency: 0.872 ± 0.08) and did not significantly differ from the latency of naïve mice (Tukey’s multiple comparisons test; *p* = 0.249). Moreover, the percentage of pups retrieved significantly decreased from 100% for all subjects on P5 to 13.33% ± 0.08 after ablation of the MOE (Figure 1D; Tukey’s multiple comparisons test, *p* = 0.002). These data are consistent with a requirement for volatile odor detection via the main olfactory epithelium for pup retrieval in surrogates.

**Figure 1:**
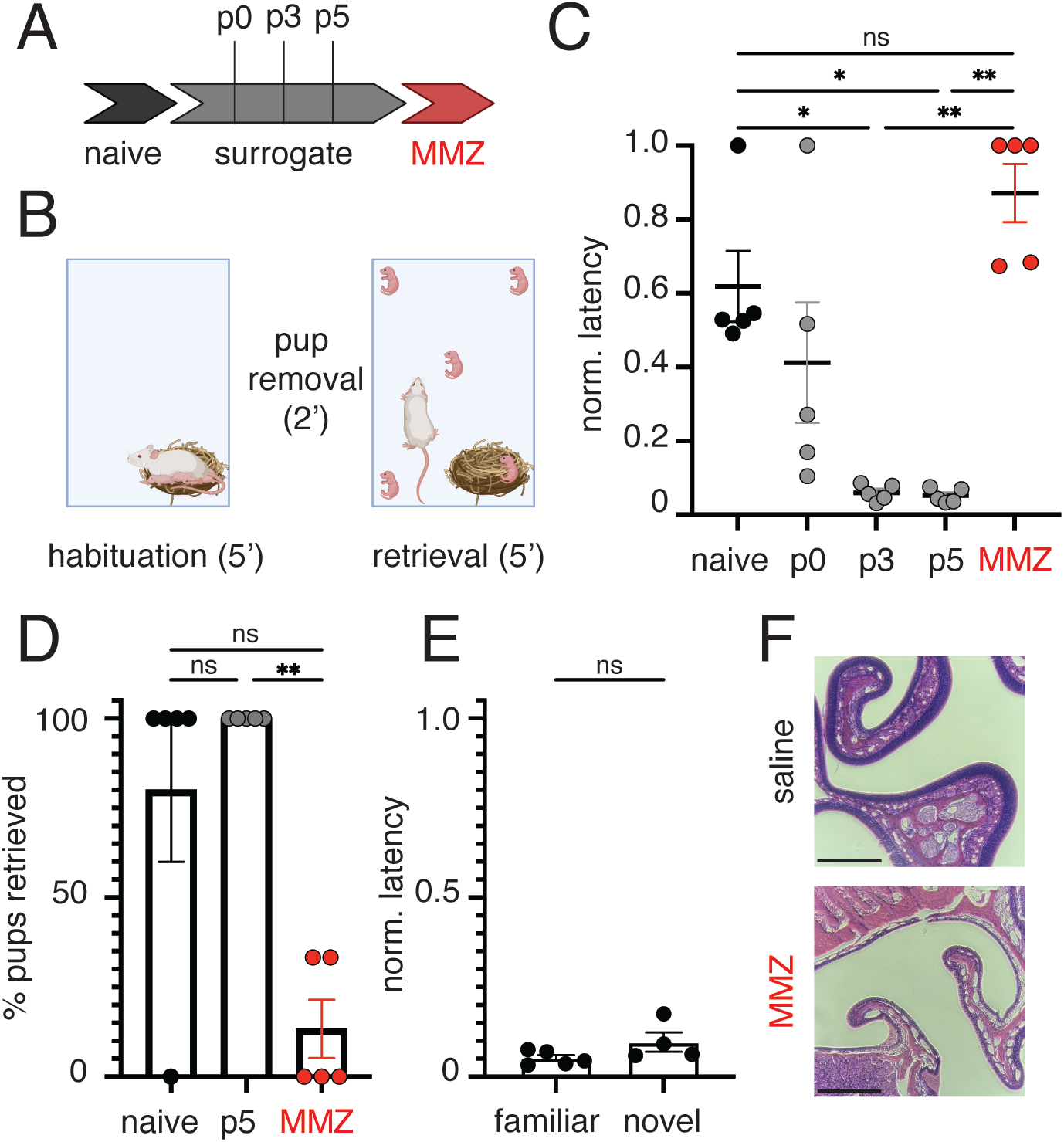
Olfactory signals from the main olfactory system are essential to maintain pup retrieval in surrogates. (A) Experimental timeline. (B) Schematic of retrieval assay. (C) Plot of latency index over time. Nulliparous female mice (n = 5) were assessed for pup retrieval as described prior to any exposure to pups (‘naïve’), on days P0, P3 and P5 relative to the birth of the pups with which they were co-housed, and again 7 days after a single IP injection of MMZ (50 mg/kg). Latency index scores (mean ± SEM) were naïve: 0.622 ± 0.0, P0: 0.413 ± 0.163, P3: 0.060 ± 0.01, P5: 0.052 ± 0.01, and MMZ: 0.872 ± 0.08. All pairwise comparisons were performed using Tukey’s multiple comparisons test (* *p* < 0.05, ** *p* < 0.01). Any comparisons not depicted were not significant (*p* > 0.05). (D) Bar plot comparing the percentage of pups retrieved among naïve mice, surrogates on P5, and surrogates after MMZ treatment. Pairwise comparisons were performed using Tukey’s multiple comparisons test (** *p* < 0.01) (E) Plot of latency index comparing pre-MMZ performance when retrieving the familiar, co-housed pups and a novel set of pups. A pairwise comparison of these values showed no significant difference (Wilcoxon matched-pairs signed rank test, *p* = 0.625). (F) Photomicrographs comparing representative paraffin-embedded, H&E stained sections of the main olfactory epithelium from a mouse that received an injection of MMZ and a mouse that received a control injection of saline (scale bar = 200 μm).

Because we allowed seven days for the MMZ treatment to complete, the original pups that were familiar to the surrogate were too large and too independently mobile for further retrieval testing, and we replaced them with a younger novel litter. We excluded the possibility that the impaired retrieval performance was simply due to the introduction of a litter of unfamiliar pups. We measured retrieval performance of the surrogates using a novel litter of pups before treatment with MMZ. The latency to retrieve novel pups and the latency to retrieve the familiar pups with which they had been co-housed did not significantly differ (Figure 1E; Wilcoxon matched-pairs signed rank test, *p* = 0.625). Based on these results, we conclude that the retrieval proficiency exhibited by the surrogates was not limited to the pups with which they were co-housed. Moreover, the poor retrieval exhibited by MMZ-treated surrogates was not due to the unfamiliarity of the litter and was more likely attributable to disrupted olfactory signaling.

### Projection neurons within the BA target the AC

Despite prior reports that pup odor can dramatically influence responses to auditory stimuli in the AC, the pathway by which odor signals reach the AC is unknown. We therefore used retrograde viral tracing methods to identify candidate inputs to the AC that could carry odor information and modulate auditory activity. We injected AAVrg-CAG-tdTomato (tdT) into the left AC as a retrograde neuronal tracer to label brain areas that project to the AC (Figure 2 A-D). In addition to known AC afferent structures such as the contralateral AC, the medial geniculate body of the thalamus (MGB), and the piriform cortex (Figure 2 C-D), we observed tdT-labeled cells in both the ipsi- and contralateral basolateral and basomedial amygdala. Henceforth, we collectively refer to these amygdala subregions as the basal amygdala (BA, Figure 2A). Neurons in the BA constitute an important component of the circuitry that governs maternal behavior.

**Figure 2:**
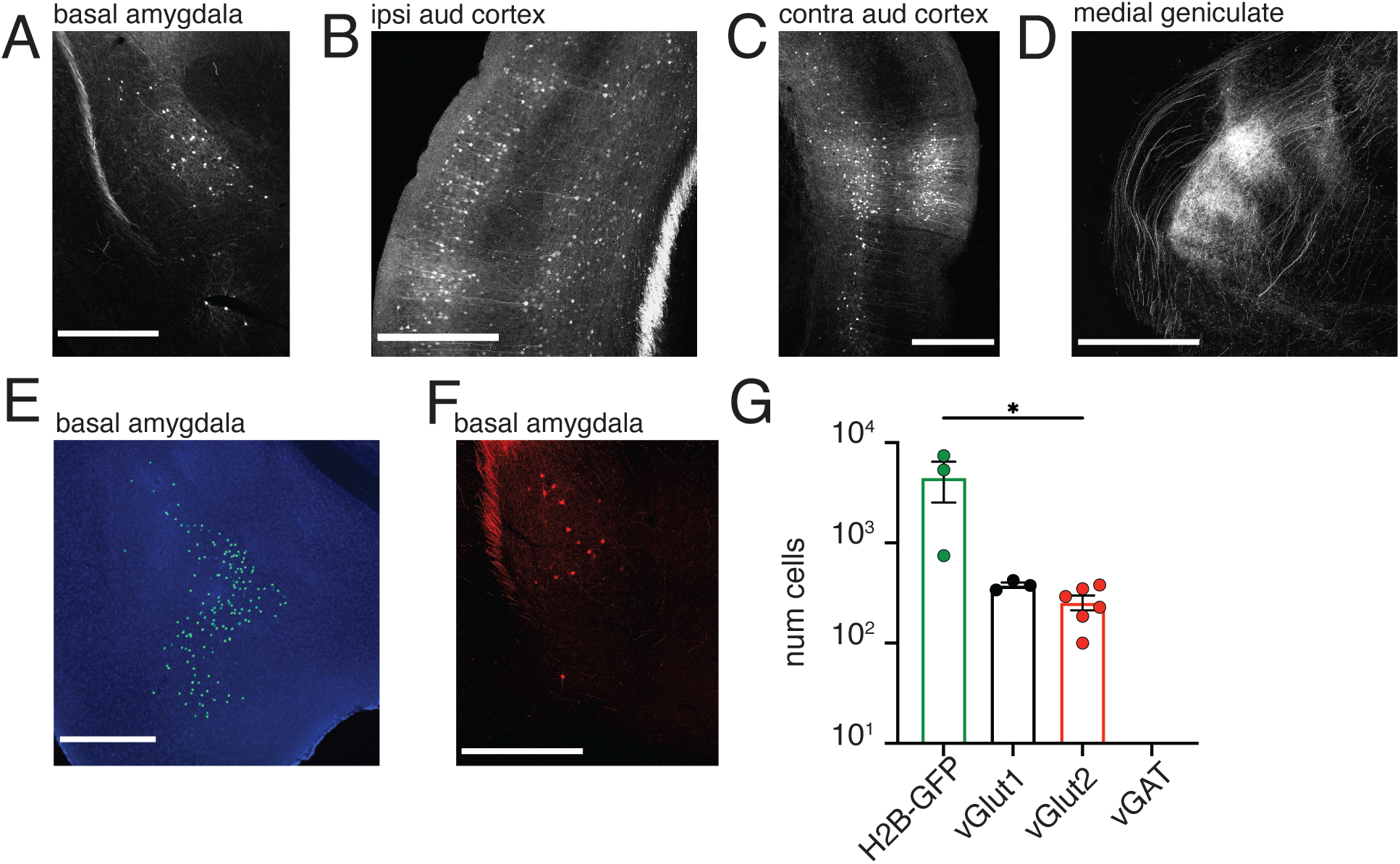
A subpopulation of glutamatergic neurons in the BA project to the AC. (A) Retrograde tracing of AAVrg-CAG-tdTomato from the AC reveals cell bodies labeled within the BA, (B) throughout the ipsilateral AC, (C) contralateral AC, and (D) medial geniculate body (scale bars = 500 μm). (E) Unrestricted retrograde tracing of AAVrg-hSyn-Cre-WPRE.hGH from the AC of a H2B-GFP mouse yields GFP positive nuclear staining for quantification of BA neurons that project to the AC (scale bars = 500 μm). (F) Cre-restricted tracing of AAVrg-FLEX-tdTomato from the AC in vGlut1Cre mouse line (scale bars = 500 μm). (G) Quantification of tracer positive neurons labeled by retrograde tracing from the AC in unrestricted (H2B-GFP, 4513 ± 1977neurons) and Cre-restricted mouse lines (vGlut1 256.3 ± 43.17 neurons; vGlut2 380.3 ± 23.99 neurons; vGAT 0 ± 0 neurons) revealed significantly fewer tracer positive neurons in vGlut2 animals compared to H2B-GFP (Mann-Whitney test, *p* = 0.0238).

They receive multimodal sensory input, including olfactory input [14], and are proposed to contribute to appetitive maternal responses by conveying sensory information to the mesolimbic reward system [15, 16]. In rats, inactivation of the BA selectively disrupts pup retrieval behavior, while minimally affecting other maternal behaviors, like nursing [17] suggesting that the BA participates in goal-directed aspects of maternal care. Moreover, neurons in the BA receive input from neighboring olfactory amygdala nuclei [18]. Based on these properties, we chose to focus our study on the population of BA neurons that project to the AC.

To quantify the extent of the BA-AC projection, we performed separate tracing experiments injecting AAVrg-hSyn-Cre-WPRE-hGH virus into the AC of Rosa26-stop^flox^-H2B-GFP mice. With this strategy, Cre activated a nuclear-localized GFP in retrogradely labeled neurons, facilitating automated cell counting. Thousands of neurons (Figure 2E; 4513 ± 1977 cells; n = 3 mice) were labeled throughout the BA. To learn more about the neurochemical identity of the neurons that contribute to the BA-AC pathway, we performed additional retrograde tracing experiments in several lines of transgenic mice that express Cre in different populations of glutamatergic or GABAergic neurons (Slc17a7^tm1.1(cre)Hze^/J, vesicular glutamate transporter type 1 (vGlut1); Slc17a6^tm2(cre)Lowl^/J, vGlut2; or Slc32a1^tm2(cre)Lowl^/J, vesicular GABA transporter (vGAT)). Not surprisingly, injections of AAV that was Cre-independent labelled the largest number of cells in the BA-AC projection neurons (Figure 2G). Injections of AAV driving Cre-dependent expression of fluorophore, AAVrg-FLEX-tdTomato resulted in less extensive labeling overall, but only in VGlut1-Cre and VGlut2-Cre mice (Figure 2F; VGlut1-Cre: 380.3 ± 23.99 neurons, n = 3 mice; VGlut2-Cre: 256.3 ± 43.17 neurons, n = 6 mice). No retrogradely-labelled neurons were found in VGAT-Cre mice (n = 3 mice). Based on our viral retrograde tracing experiments, we conclude that a relatively large, but heterogeneous population of glutamatergic excitatory neurons in the BA project to the AC (Figure 2G).

### BA-AC projection neurons respond to odors including pup odor

To determine whether the BA-AC projection neurons were sensitive to pup-related odors, we used fiber photometry in awake head-fixed mice to measure bulk Ca^2+^signals from BA-AC neurons in response to olfactory stimuli. We selectively labeled the BA-AC projection neurons by using the intersectional viral strategy depicted in Figure 3A (see also Figure S1). Briefly, we injected retrograde AAVrg-hSyn-Cre-WPRE-hGH into the AC and made an injection of AAV5-syn-FLEX-GCaMP6s-WPRE into the BA, thereby expressing the fluorescent Ca^2+^ sensor GCaMP6s exclusively in BA neurons that project to the AC. After three weeks to allow for viral expression, naïve virgin females were habituated for several days to head fixation on a wheel and then presented with pseudorandomized two second (s) trials of monomolecular odorants or pup odor and clean nesting material every 30 s (Figure 3B).

**Figure 3:**
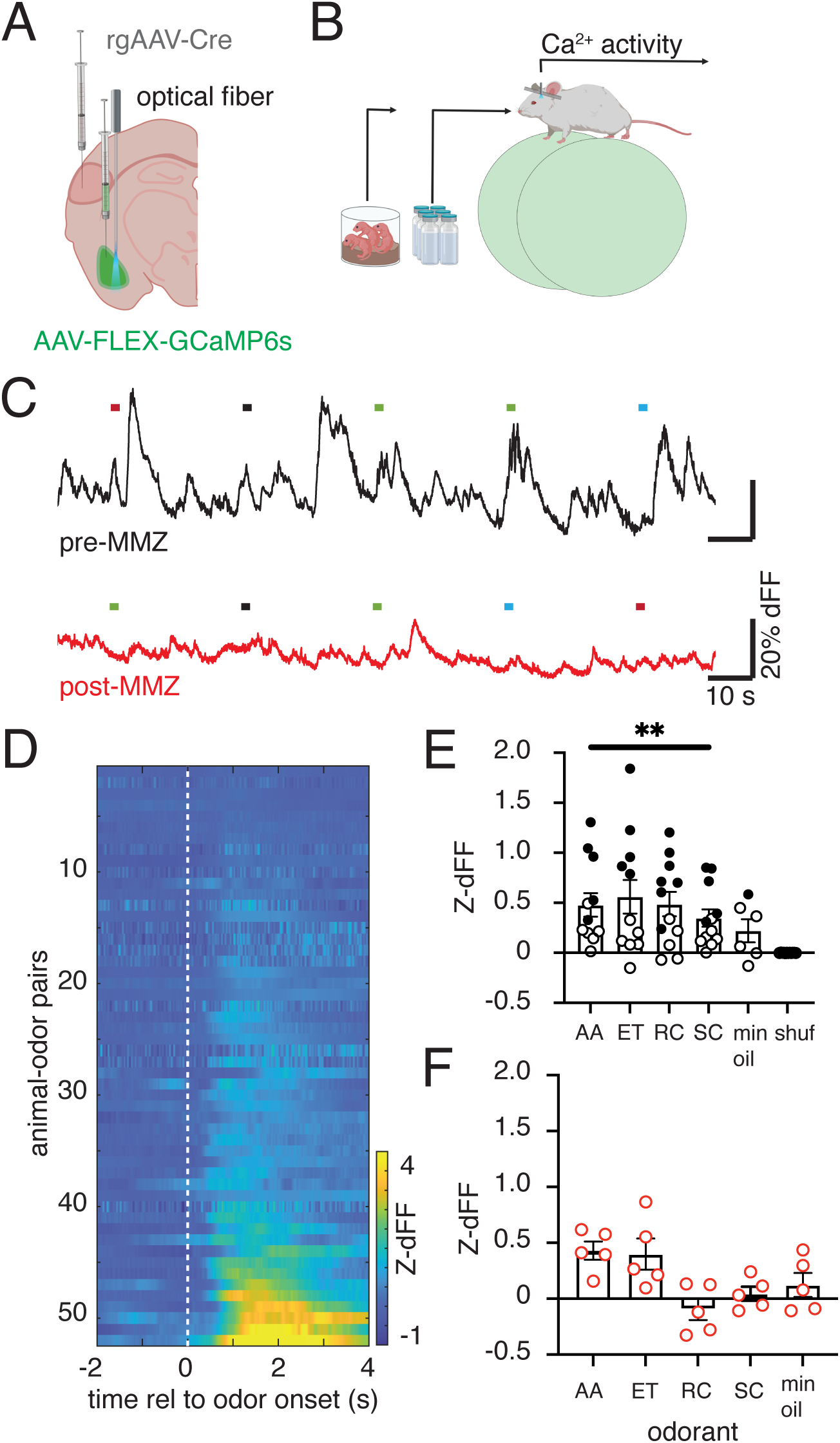
BA-AC projection neurons exhibit increased Ca^2+^ activity in response to olfactory stimuli. (A) Schematic of our intersectional viral strategy for selectively labeling BA-AC projection neurons with GCaMP6s and recording ongoing Ca^2+^ activity (B) Schematic of head-fixed recording of Ca^2+^ activity during presentation of monomolecular odorants or pup odor. (C) Raw dFF traces from one animal during the presentation of monomolecular odorants before MMZ treatment (top panel, black trace) and after MMZ treatment (bottom panel, red trace). Dashed lines are color-coded by odor stimulus. Black dashes are mineral oil controls. (D) Heatplot of mean responses to all odors by all mice. Each row in the plot represents the mean response over multiple trials to one odor in one mouse (odor-mouse pair), expressed as a Z score (Z-dFF), according to the color bar. Rows are ordered by the magnitude of each response. (E) Quantification of the increased Ca^2+^ signal during odor presentation. (top panel): Responses to all odors were observed among recorded subjects (n = 13 mice) Filled circles indicate mean responses that were outside the center 95% of a dummy distribution generated with a bootstrap procedure using data from the same mouse (see Methods). For each odor, the responses of all mice were compared to a mean response of 0: amyl-acetate (AA: 0.4788 ± 0.1180), S-carvone (SC, 0.3478 ± 0.0859), ethyl tiglate (ET: 0.5605 ± 0.186), R-carvone (RC: 0.4868 ± 0.1220), linalool (LN: 0.6282 ± 0.2049) mineral oil (min oil, 0.2215 ± 0.1159), and shuffled data (shuf, - 0.0002 ± 0.0005). Wilcoxon signed rank test, \*\**p* < 0.01 (bottom panel). (F) No significant odor responses were observed to any odor, at either the population or individual mouse level following treatment with MMZ (AA: 0.3679 ± 0.1182; ET: 0.2452 ± 0.1338; RC: 0.1408 ± 1090; SC: −0.0559 ± 0.1402, min oil: 0.1916 ± 0.0631, n = 5).

We observed robust, odor-evoked changes in fluorescence in response to monomolecular odors (Figure 3C-E). Most odors in most mice elicited an abrupt increase in GCaMP6s fluorescence, although the magnitude of Ca^2+^ signals was variable depending on the individual subject and the odor presented (Figure 3D and Figure S2). Comparing the stimulus-evoked signal to baseline for all mice, mean Ca^2+^ fluorescence showed a significant increase for each of four monomolecular odors presented (Figure 3E). We confirmed that the responses to these stimuli were mediated by the main olfactory system by ablating the MOE with MMZ. After ablation, we no longer observed significant responses to any of the monomolecular odors (Figure 3 F).

In addition to the arbitrary, non-social odors, we presented pup odor by collecting the headspace of a glass jar containing several pups (age 2 – 4 days) and directing the stream of odorized air onto the nose of a head-fixed, fiber-implanted mouse. In virgin females without maternal experience (“naive”), we observed a clear increase in Ca^2+^ signal in response to pup odor that had a similar temporal structure to the responses to non-social odors (Figure 4 A-B). Pup odor evoked responses persisted throughout the surrogacy experience (Figure 4 B-C, E). As seen with other odors, this response was abolished after MMZ treatment (Figure 4 C-D). No odor-evoked changes in fluorescence were observed in control animals injected to express GFP in BA-AC neurons instead of GCaMP6s (Figure S3A; paired t-test comparing Z-dFF at baseline (2 s) to after odor delivery (3 s), *p* > 0.05, Figure S3B; Wilcoxon signed rank test, *p* > 0.05), verifying that the fluctuations we measured in response to odors reflect neural activity and not movement or other artifacts. Based on the above data, we conclude that the BA-AC projection neurons respond to pup odor and therefore constitute a likely candidate pathway for odor information to access the AC.

**Figure 4:**
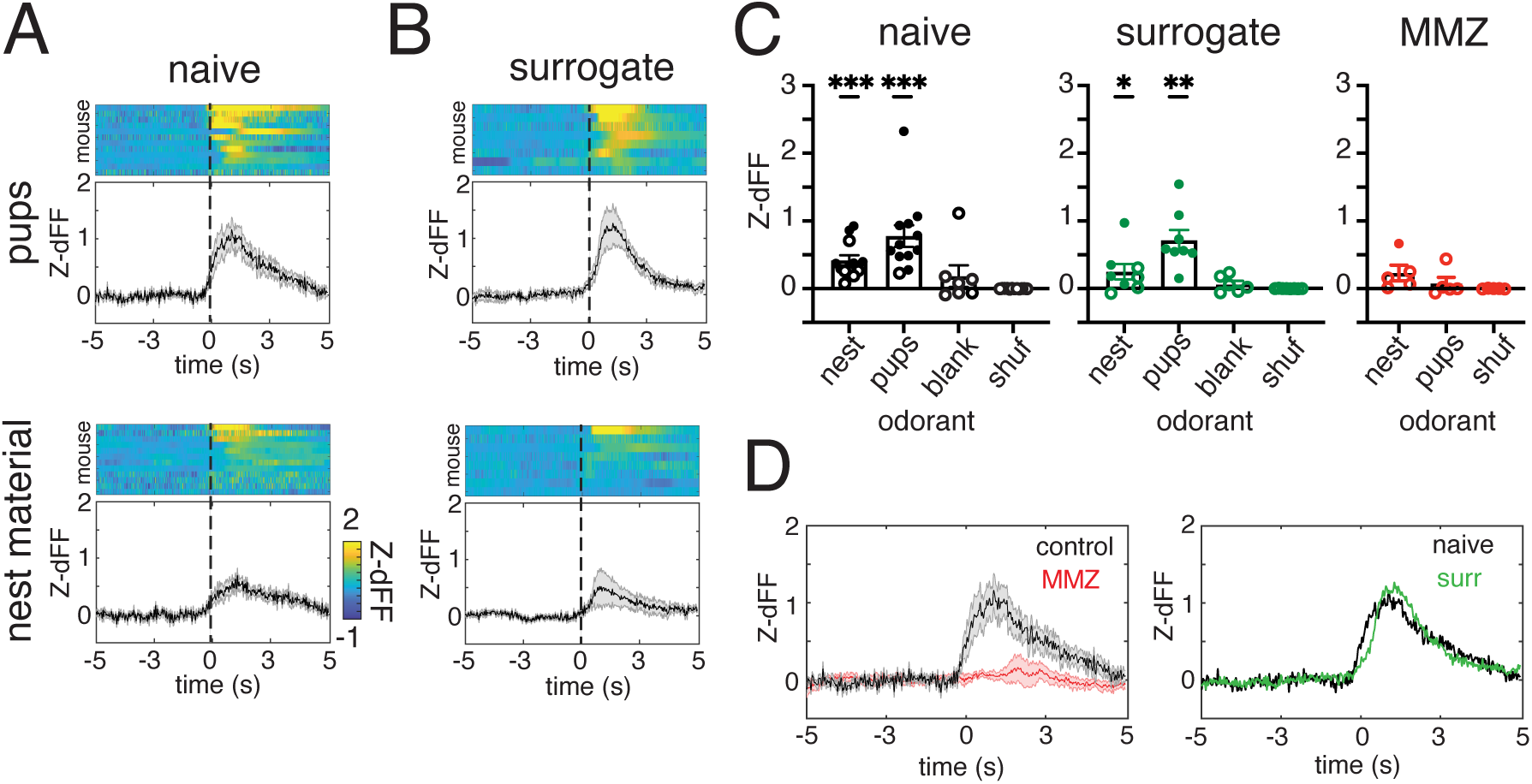
Pup-odor-evoked activity is maintained across surrogacy. (A) Normalized evoked responses to pups and clean nest material in naive and (B) surrogate animals. Heatmaps were sorted by decreasing strength of response (top panels) to demonstrate the variability of average odor-evoked activity across animals with SEM indicated by shaded area (bottom panels). Color scale applies to all heatmap plots. (C) Quantification of the Z-dFF signal during odor presentation in naive animals (nest: 0.4159 ± 0.0778, pups: 0,7764 ± 0.1597, blank: 0.1844 ± 0.1603, and shuffled data: −0.0005 ± 0.0011, n = 7-12, closed circles indicate mean responses that exceeded a 95% cutoff of bootstrapped data), surrogates (nest: 0.2487 ± 0.1147, pups: 0.7158 ± 0.1492, blank: 0.0655 ± 0.0489, and shuffled data: −0.0001 ±0.0001, n = 6-8), and after MMZ treatment (nest: 0.2301 ± 0.1164, pups: 0.0766 ± 0.0912, and shuffled data: −0.0001 ± 0.0025, n = 5). Each point represents a single animal. Bars indicate mean ± SEM Wilcoxon signed rank test. (D) Average pup-odor evoked activity comparing response from naive animals (black trace, n = 12) and MMZ treated animals (red trace, n = 5), and (E) naive (black trace, n = 12) versus surrogate animals (green trace, n = 8).

The property of responding to pup odor was consistent among all mice when recording from BA-AC neurons. In addition to neurons that project to the auditory cortex, BA contains separate neurons that project to either nucleus accumbens (NAcc), the ventral hippocampus, or other targets. To assess whether response to volatile odors from pups is a shared property among all neurons in BA, we performed head-fixed recordings from BA neurons that project to the NAcc (BA-NAcc) while presenting clean nesting material, chow, and pup odor. Our viral strategy for labeling BA-NAcc neurons was very similar to the strategy we used to label the BA-AC (Figure 3A). In this case, we expressed GcaMP6s in BA-NAcc by injecting retrograde AAVrg-hSyn-Cre-WPRE-hGH into NAcc and AAV5-syn-FLEX-GcaMP6s-WPRE into the BA (Figure S4A).

After ∼three weeks to allow for expression, nulliparous adult female mice (n = 6), were delivered a two s trial of either pup odor, chow, or clean nesting material in pseudorandomized order every 30 s (Figure S4B, C). All mice consistently exhibited no response to pups or the blank (empty vial) stimulus; several mice exhibited responses to nest material and chow, but when comparing mean odor-evoked responses of all mice to each odor with zero, only responses to chow achieved statistical significance (n = 6, one sample t-test; *p* < 0.05). These results demonstrate that information regarding pup odors is shared with AC, but not all BA targets.

### BA-AC neurons are active during pup-seeking

Having discovered that BA-AC neurons were sensitive to pup odor, we next asked how they may participate in free interactions with pups, including retrieval. To do this, we recorded Ca^2+^ activity from BA-AC neurons in freely behaving females daily as they performed pup retrieval from the naïve state through surrogacy. Based on the results of the head-fixed experiments, we hypothesized that we would observe odor-evoked responses in the BA-AC neurons when a naïve female engaged with pups by picking them up or sniffing them during the retrieval behavior.

Surprisingly, by aligning the fluorescent traces to the mouse’s behavior, we observed that while searching for pups (defined here as the time between egress from the nest and either contact with a pup or returning to the nest), Ca^2+^ signals were elevated (Figure 5A). For a more systematic analysis, we segmented pup retrieval into four behavioral events: pup retrieval, pup sniffing, air sniffing and search. When GCaMP6s traces were aligned to each phase of retrieval, BA-AC neurons only showed a significant increase in activity as the females (n = 10 mice) searched the cage for a pup (Figure 5D; Wilcoxon signed rank test, *p* = 0.027), not when they made contact to retrieve or engaged with a pup physically by sniffing it. In fact, the fluorescent signal dropped significantly as soon as contact with a pup was made for retrieval (Figure 5B-D, Wilcoxon signed rank test, *p* = 0.004) or investigation (Figure 5D; Wilcoxon signed rank test, *p* = 0.010). In Figure 5B, the variability seen among the individual trials, and the ramping appearance leading up to contact are due to the fact that animals can spend very different amounts of time on individual search bouts. These results suggest that the BA-AC circuit is primarily active during exploratory and/or goal-directed aspects of maternal behavior.

**Figure 5:**
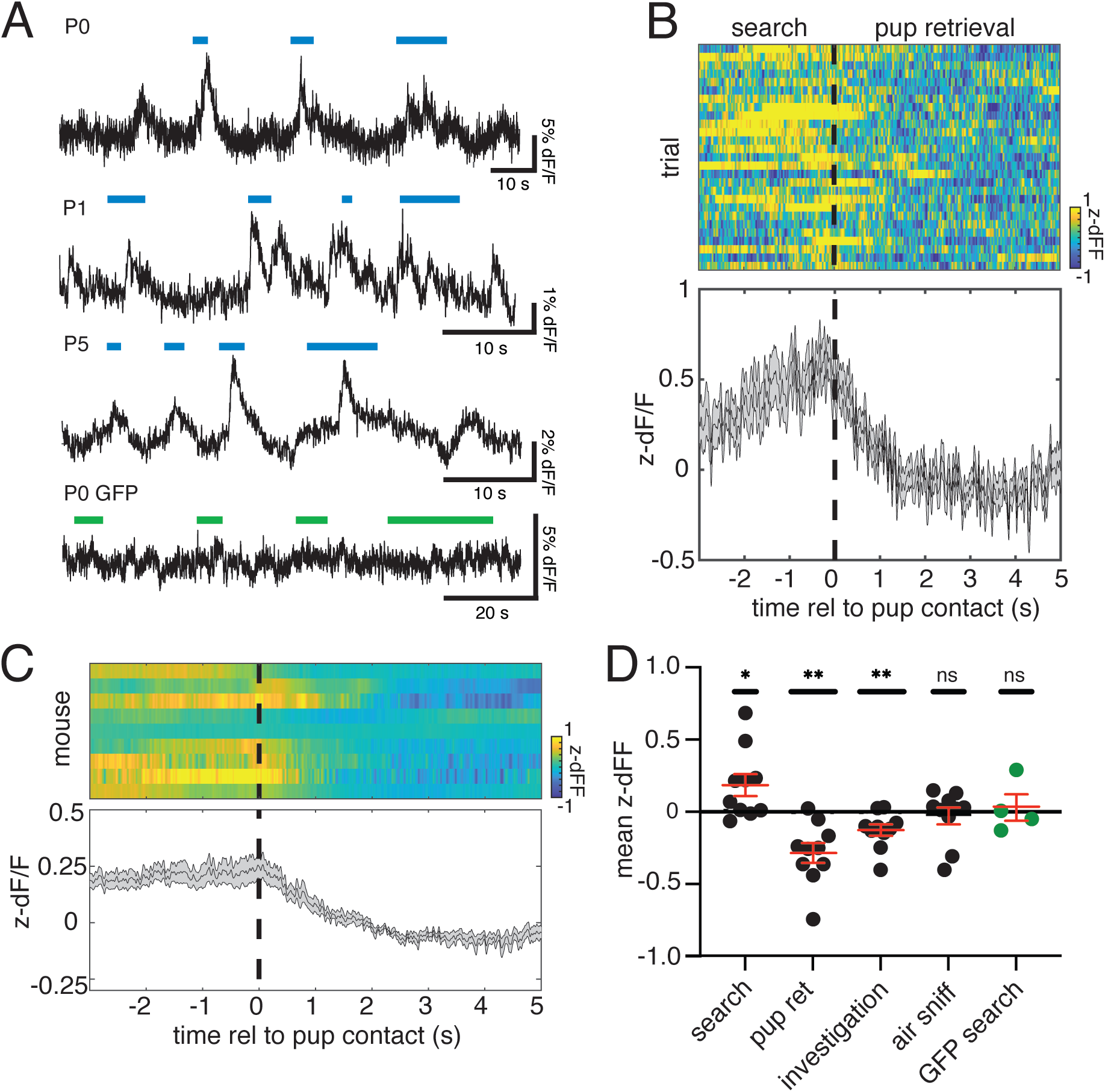
BA-AC neurons show elevated activity during pup search that terminates with pup contact. (A) Example traces of dFF during several episodes of searching for pups. Each trace is taken from a different mouse on a different postnatal day (shown to the upper left of each trace). The colored bars above each trace denote time spent searching for a pup, beginning at the time the female exits the nest and ending at the time she contacts a pup. The blue bars are above traces taken from mice expressing GCaMP6s in BA-AC neurons. The trace with the green bars above it was taken from a mouse that was expressing GFP in BA-AC neurons. (B) Average data across all retrieval trials from one mouse aligned to the end of the search period. The upper panel is a heat map of Z-dFF for 27 retrieval events taken from P0, P1, P3, and P5. Each row depicts one retrieval event aligned to the end of the search period, when the mouse contacts the pup (vertical dashed line). Color is mapped to Z-dFF according to the color bar on the lower right. The lower panel is a plot of mean Z-dFF for all trials. The shaded area around the trace denotes the SEM of the response. Note the abrupt decrease in activity as the female encounters the pup. (C) Plot summarizing the average response to pup contact in all mice. The upper panel is a heatmap of the average response to a search that terminates in an encounter with a pup across all trials for all mice (n = 9). Each row is the mean response to pup contact (vertical dashed line) for one mouse, calculated as in (B). Color is mapped to Z-dFF according to the color bar on the lower right. The lower panel is a plot of mean Z-dFF for all mice. The shaded area around the trace denotes the SEM of the response. (D) Plot of the mean change in fluorescence associated with several events (search, pup contact, investigation, and air sniff) across all mice. Each point denotes the mean response of one mouse, defined as the difference between the mean fluorescence during the first three s after each event and the mean value during the immediately preceding three s. Mean activity during search (0.1863 ± 0.076 Z-dFF), after pup contact (−0.284 ± 0.068 Z-dFF), and during investigation (−0.126 ± 0.040 Z-dFF) were significantly different from 0 (n = 10 mice; Wilcoxon signed rank test, \**p* < 0.05, \*\**p* < 0.01). No significant change in fluorescence was detected when the mouse sniffed the air (n = 10 mice; −0.028 ± 0.057 Z-dFF, Wilcoxon signed rank test, *p* = 0.695) or in mice that expressed activity-independent GFP in the BA (n = 4; 0.030 ± 0.092 Z-dFF, Wilcoxon signed rank test, *p* > 0.999).

### Optogenetic activation of BA-AC neurons bidirectionally modulates AC activity

Acutely presenting pup odor modifies auditory responses in the AC of female mice with maternal experience. Having established that the BA-AC projection neurons are responsive to pup odor, and are active while the female searches for pups, we therefore hypothesized that selective optogenetic activation of the BA-AC pathway may modulate auditory responses in AC. We tested this hypothesis using an intersectional viral strategy to selectively label the BA-AC projection neurons with either the excitatory opsin, Channelrhodopsin-2 (ChR2), or GFP as a control (Figure 6A). After several weeks to allow for stable viral expression, mice in both groups were used for acutely anesthetized extracellular recording experiments. Auditory cortical neurons were recorded with the ‘loose patch’ technique [5, 19] while presenting a set of tones and pup vocalizations (calls) in pseudorandom order (see Methods). On 50% of trials, BA-AC axon terminals labelled with ChR2 were activated by directing blue light (473nm) onto the cortical surface (Figure 6B). To examine the effect of maternal experience, recordings were performed in either naïve virgins or surrogate mothers.

**Figure 6:**
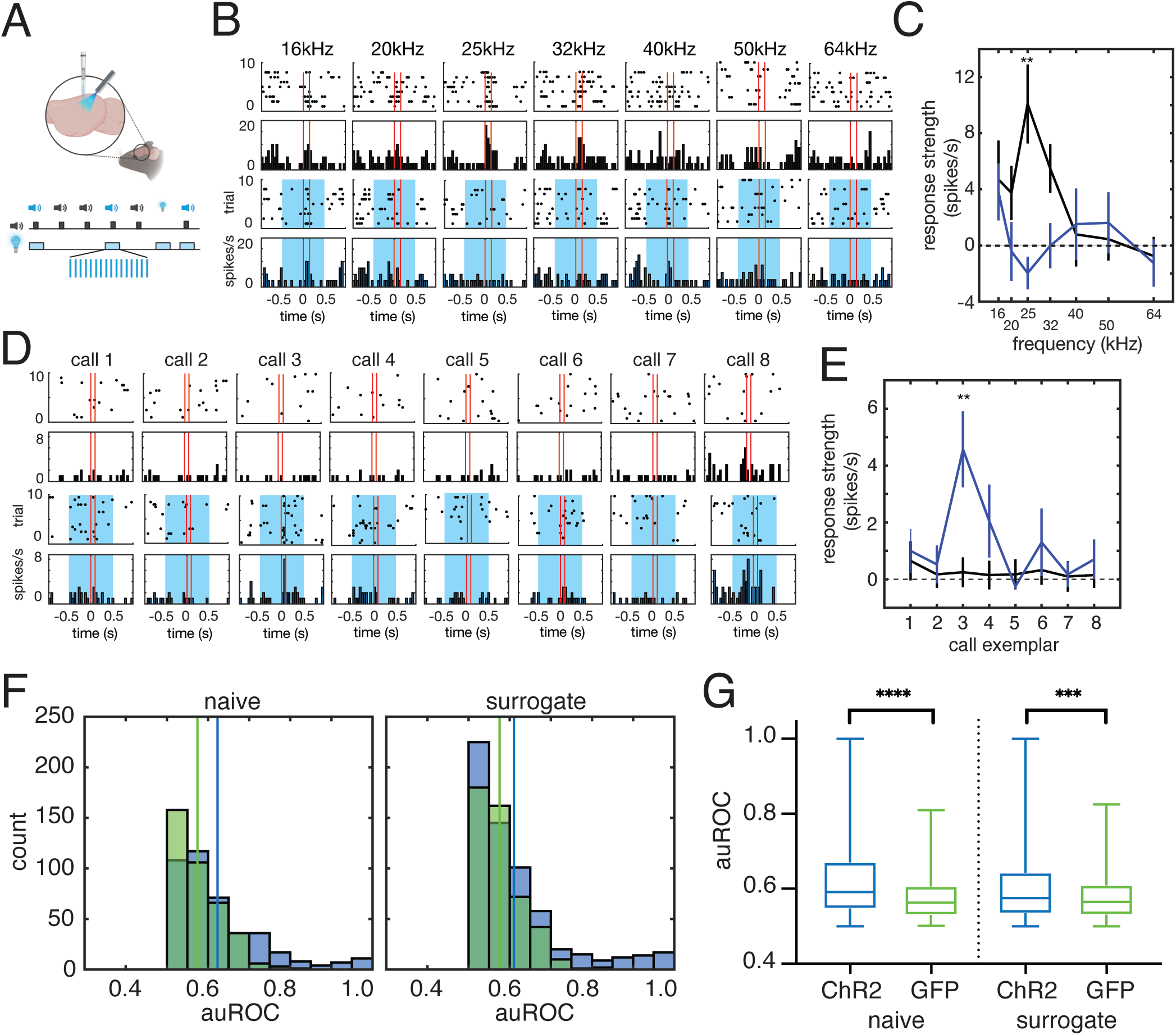
Optogenetic activation of the BA-AC pathway elicits widespread and bidirectional modulation of auditory responses in the auditory cortex. (A) Schematic of the experimental design. The same intersectional strategy for labeling BA-AC neurons with GCaMP6s (Figure 3A) was used to express ChR2 in the same neurons. The retrograde virus AAVrg-hSyn-Cre-WPRE-hGH was injected into the AC and at the same time, the Cre-dependent AAV9-CAGS-FLEX-ChR2-tdT-WPRE-SV40 was injected into the BA. Mice were acutely anesthetized for electrophysiology recordings and presented with auditory stimuli including synthesized pure tones and previously recorded pup calls. Stimuli were presented in a pseudorandom order and 50% of trials for each stimulus were accompanied by a train of 473 nm light pulses (20 Hz) directed at the cortical surface to activate BA-AC terminals. (B) Plots comparing responses of an auditory cortical neuron to logarithmically spaced pure tones when presented during light activation of BA-AC and when presented alone. Data are from a naïve female. Each stimulus is associated with a raster plot and peristimulus time histogram (bin size = 50 ms) from control trials (top row) and from light activation trials (bottom row with blue shading). The blue shading denotes the duration of the light train relative to the tone. Note that responses to the 25 kHz tone are significantly weaker when BA-AC terminals were activated by light (n = 8 trials; comparison of trials with and without light, unpaired t test with Bonferroni correction, \*\**p* < 0.01). (C) Line plot of data from (B) comparing the mean baseline-subtracted firing rate evoked by each tone on control trials (black) and on light trials (blue) (\*\**p* < 0.01). Vertical lines denote SEM. (D) Plots comparing responses of a different auditory cortical neuron to 8 different pup call exemplars when presented during light activation of BA-AC and when presented alone. Data are from a surrogate female. Panels are organized as in (B). (E) Line plot of data from (D) comparing the mean baseline-subtracted firing rate evoked by each call on control trials (black) and on light trials (blue). Note that in this case, responses to call 3 are significantly stronger when BA-AC terminals were activated by light (n = 8 trials; comparison of trials with and without light, unpaired t-test with Bonferroni correction, \*\**p* < 0.01). (F) Distribution of auROC values for each cell-stimulus pair from ChR2 injected animals (blue) and GFP controls (green) separated by pup experience (naïve, left panel; surrogate, right panel). The mean value of each distribution is marked by the vertical line of the corresponding color. (G) Box plots of the data in (F) demonstrate that ChR2 expressing animals exhibit significantly greater discriminability between light-on and light-off trials for both naïve virgins (ChR2: n = 415 cell-stimulus pairs, 0.623 ± 0.005; GFP: n = 376 cell-stimulus pairs, 0.574 ± 0.003, Mann-Whitney U-test, *p* < 0.0001) and surrogates (ChR2: n = 616 cell-stimulus pairs, 0.610 ± 0.005; GFP: n = 469 cell-stimulus pairs, 0.576 ± 0.003, Mann-Whitney U-test *p* = 0.0005).

We recorded 83 neurons from 17 ChR2 injected mice (34 from 7 naïve mice and 49 from 10 surrogates) and 72 units from 15 GFP injected mice (33 from 7 naïve and 39 from 8 surrogates). All neurons exhibited stimulus-specific activity, and in ChR2-expressing mice, responses to some or all stimuli were modified by optogenetic activation. For example, on trials without light stimulation, the neuron depicted in Figure 6B-C showed significant increases in firing to 16, 20, 25, and 32 kHz tones. However, in trials with light activation of the BA-AC terminals, responses to several tones were abolished (Figure 6B, lower panels and Figure 6C, blue trace). In some neurons, stimuli that elicited no response on trials without optogenetic activation, actually did evoke a significant response on trials with light (Figure 6D, E).

We quantified the magnitude and the prevalence of light modulation for all neurons and stimuli by performing a receiver operator characteristic (ROC) analysis for each cell-stimulus pair (see Materials and Methods). In this analysis, the area under the ROC curve (auROC) can be used as a measure of the discriminability between light-on versus light-off trials. We found that in naïve virgins, the auROC values for units recorded from ChR2 animals were significantly greater than those observed in GFP controls (Figure 6F-G; ChR2: 0.62 ± 0.01; GFP: 0.574 ± 0.00, Mann-Whitney U-test, *p* < 0.0001). The same was true in surrogates (Figure 6F-G, ChR2: 0.61 ± 0.01; GFP: 0.58 ± 0.00, Mann-Whitney U-test p = 0.0005). Therefore, the modulation of auditory responses was due to optogenetic activation and not light itself.

The direction of modulation was not accounted for in the above analysis, but it varied by neuron and stimulus. We observed examples of both optogenetic enhancement *and* suppression of auditory response strength in naïve females (Figure 6C, E) and surrogates (Figure S5).

However, a comparison of the effect of optogenetic activation of the BA-AC pathway revealed that maternal experience flipped the balance from predominantly suppression in naïve females (Figure 7A) to predominantly enhancement in surrogates (Figure 7B).

**Figure 7:**
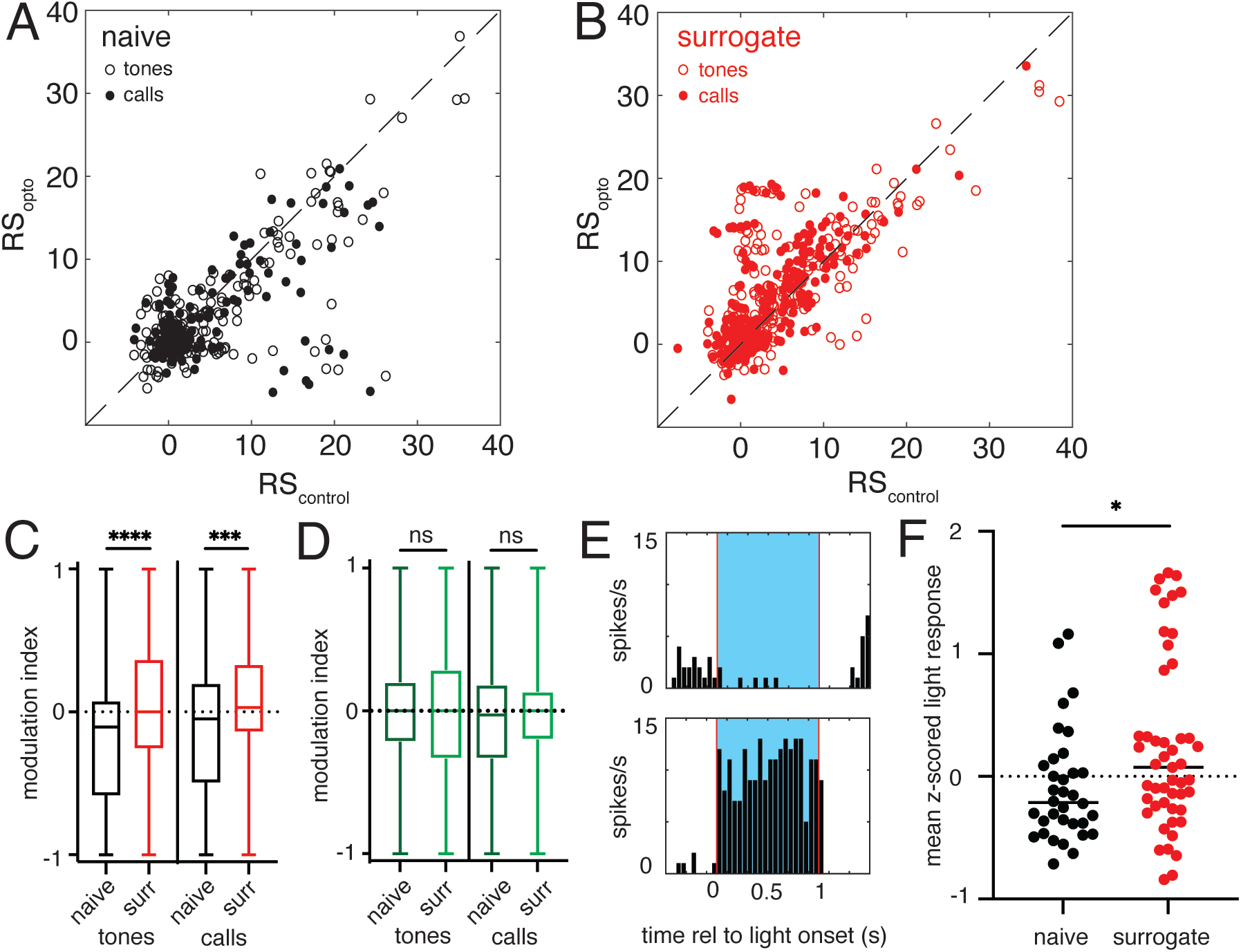
The effects of BA-AC activation on AC neurons are experience-dependent. (A) Scatterplot of response strength (RS) comparing control trials versus light activation trials for all cell-stimulus pairs in naïve mice expressing ChR2 in BA-AC neurons. Tone stimuli are plotted as open circles and call stimuli are plotted as filled circles. (B) Scatterplot of response strength (RS) comparing control trials versus light activation trials for all cell-stimulus pairs in surrogate mice expressing ChR2 in BA-AC neurons. Tone stimuli are plotted as open circles and call stimuli are plotted as filled circles. (C) Box plot of modulation index comparing modulation of tone responses in naïves (black) and surrogates (red) expressing ChR2 in BA-AC (naïve: n = 226 cell-stimulus pairs, −0.185 ± 0.037; surrogates: n = 291 cell-stimulus pairs, 0.031 ± 0.034; Mann-Whitney U test, *p* < 0.0001) (left side). On the right side, the box plot compares modulation of USV call responses in naives (black) and surrogates (red) (naïve: n = 175 cell-stimulus pairs, − 0.110 ± 0.042, surrogate: n = 274 cell-stimulus pairs, 0.055 ± 0.032; Mann-Whitney U test, *p* = 0.0006 Mann-Whitney U test). (D) Box plot of modulation index comparing modulation of tone responses in naïves (dark green) and surrogates (light green) expressing GFP in BA-AC (naïve: n = 213 cell-stimulus pairs, −0.027 ± 0.031; surrogates: n = 223 cell-stimulus pairs, −0.005 ± 0.038; Mann-Whitney U test, *p* = 0.453) (left side). On the right side, the boxplot compares modulation of USV call responses in naives (black) and surrogates (red) (naïve: n = 153 cell-stimulus pairs, −0.078 ± 0.040, surrogate: n = 206 cell-stimulus pairs, −0.020 ± 0.030; Mann-Whitney U test, *p* = 0.204). (E) Example responses to light only trials for naïve and surrogate mice. The top panel shows a peristimulus time histogram (bin size = 50 ms) of the mean firing rate of an AC neuron in a naive female during 20 trials of light stimulation accompanied by silence. The bottom panel shows a peristimulus time histogram (bin size = 50 ms) of the mean firing rate of an AC neuron in a surrogate female during 20 trials of light stimulation accompanied by silence. (F) Swarm plot comparing the mean response to light only trials for all neurons in ChR2-expressing mice. (naïve: n = 34, −0.091 ± 0.089; surrogates: n = 49, 0.237 ± 0.101, Mann-Whitney U, *p* = 0.0248).

To determine whether stimulus frequency might influence this categorical outcome, we used Fisher’s exact test to compare the number of units enhanced and suppressed by optogenetic stimulation from ChR2 injected animals versus GFP controls. For the majority of tones presented, the effect of optogenetic activation on AC responses to sound was independent of tone frequency (Figure S6A). The only exception was naïve responses to 32kHz tones (Fisher’s exact test, *p* = 0.02). Similarly, the effect of activation of the BA-AC pathway was independent of the call exemplar presented, with the exception of surrogate responses to call exemplar 5 (Figure S6B, Fisher’s exact test, *p* = 0.0003).

To quantify the relative preponderance of optogenetic enhancement vs. suppression observed in surrogates, we calculated a modulation index (see Methods) to measure the change in response strength between light off and light on trials. We observed significant differences in modulation index between the naïve and surrogate animals for both tone stimuli (Figure 7C; naïve: n = 226 cell-stimulus pairs, −0.19 ± 0.04; surrogate: n = 291 cell-stimulus pairs, 0.031 ± 0.03; Mann-Whitney U test, *p* < 0.0001) and USVs (Figure 7C; naïve: n = 175 cell-stimulus pairs, −0.110 ± 0.042, surrogate: n = 274 cell-stimulus pairs, 0.055 ± 0.032; Mann-Whitney U test, *p* = 0.0006). There was no difference in the modulation index between naïve and surrogate GFP control mice for either tone or USV stimuli (Figure 7D; Mann-Whitney U test, p ≥ 0.05). Similar to what we observed in the number of enhanced units, the distribution of modulation indices was independent of stimulus identity (Figure S6C-D; naïve, 2-way ANOVA, p > 0.05, n = 34 units, 7 mice; surrogates, 2-way ANOVA, p > 0.05, n = 46 units, 10 mice).

In some cases, we observed a change in firing rate on light only trials, during which no auditory stimulus was played (e.g. Figure 7E). While only some cells exhibited a change in firing on light only trials, considering all cells from ChR2-expressing mice, we found that light only responses were significantly more positive in surrogates as compared to the effect of optogenetic BA-AC activation in naïve mice (Figure 7F; naïve: n = 34 neurons, −0.09 ± 0.08; surrogates: n = 49 neurons, 0.24 ± 0.10, Mann-Whitney U test, *p* = 0.025). Again, we discovered a significant experience-dependent bias in the strength of the light only response. In naïve animals, ongoing activity was suppressed by activation of the BA-AC pathway, while in surrogates it was enhanced.

## DISCUSSION

Like most social encounters, interactions between rodent pups and their parents are multisensory. They include somatosensory components and, most relevant to this work, auditory and olfactory components. The importance of pup vocalizations for eliciting retrieval is clear from several observations. First, dams will phonotax towards playback of natural USVs and synthetic USVs within the appropriate high-frequency range [20]. Second, pups that are congenitally mute are largely ignored by their mothers, even when they are separated from the nest [21]. Third, many studies have shown that initial maternal experience in both dams and virgin surrogates is correlated with plasticity in auditory responses in the auditory cortex (AC) [5–9]. Finally, in our prior work on the impairment of maternal behavior in a mouse model of Rett syndrome (*Mecp2^het^*), we showed that knocking out *Mecp2* only in the AC post-development was sufficient to substantially impair retrieval performance [6, 22].

Although less well-studied, olfactory cues from the pups are also acknowledged to be essential for pup retrieval. For example, interfering with a mother’s ability to detect volatile pup odors, either by genetically inactivating crucial components of olfactory signaling or by washing the pups to remove their odor, significantly diminishes retrieval performance [8, 11–13]. Notably, here we confirmed and extended this finding to include virgin surrogates that had already established retrieval proficiency. By using the tissue-specific toxicant MMZ to ablate the main olfactory epithelium (MOE), we demonstrated that they almost completely lost the ability to gather pups. This raises several possibilities for the role olfactory cues play in maternal retrieval behavior: they may help guide the female to the location of the pup, they may influence the perception of USVs, and/or they may trigger maternal experience-induced plasticity in the AC.

Despite the shared importance of USVs and pup odor for maternal care, relatively little is known about the circuits that coordinate audition and olfaction during maternal behavior. Previous work by Cohen and colleagues [8, 9] showed that the odor of pups directly interacts with AC responses to sounds, including USVs. Importantly, this interaction was only observed in mothers or surrogates with maternal experience. However, the circuit by which the pup odor accessed the AC was not identified.

Here we attempted to identify a specific neural pathway that could integrate the influence of odor with sensory representations of USVs in the AC. We began by injecting viral tracers into the AC to reveal candidate inputs. We identified the basal amygdala (BA) as a region that could carry information regarding odors due to its connections to olfactory areas of the amygdala [18]. We also observed sparse labeling of neurons in the piriform (olfactory) cortex. Nevertheless, in light of prior studies linking the BA with maternal behavior (see below), we instead chose to focus on its projection to the AC.

We made the following observations regarding the BA-AC pathway: First, we identified the BA neurons that project to the AC as likely predominantly glutamatergic. Second, with fiber photometric recordings exclusively from BA neurons that project to the AC, we demonstrated that these neurons robustly respond to pup odor and other odors more broadly. Third, we found that BA-AC neurons are active in freely behaving surrogates, particularly while the mouse is searching for pups, and rapidly become less active when she contacts them. Fourth, we found that optogenetic activation of the terminals of BA-AC projections changed the response of AC neurons to sound, sometimes dramatically. Finally, in naïve virgins, activation predominantly led to inhibition of auditory responses, while in surrogate mice, activation predominantly led to excitation. We conclude that a glutamatergic projection from the basal amygdala regulates auditory cortical activity in an experience-dependent manner during pup interactions, in part driven by odors.

### Implications for the auditory cortex

The pathway from the BA to the AC has not been widely studied. Responses to sound in AC are modulated by emotionally-charged stimuli [23–28], but evidence that the modulation comes directly from the amygdala is sparse. We hypothesize that the BA-AC pathway is a potential conduit for information about valence to reach AC. This may have important short-term and long-term consequences for how the AC responds to behaviorally-significant sounds. Cohen et al. [8] reported a significant influence of pup odor on responses to a variety of sounds. However, the design of their experiments precluded a trial-by-trial assessment of this influence. Instead, they recorded responses in alternating blocks lasting tens of minutes either with or without pup odor. We used an optogenetic approach that enabled precise optical control of BA inputs on the temporal scale of individual sounds. These experiments demonstrate that the BA projection is capable of rapidly modulating auditory responses. On the other hand, optogenetic activation exhibited long-term experience dependence. Modulation of the AC by pup odor was only observed in mice with pup experience, not naïve virgin mice. In our experiments, pup odor responses in the BA (at least in head-fixed surrogates) were evident prior to pup exposure and continued at least until P5. Moreover, after pup exposure, the effects of activating BA inputs shifted from being predominantly inhibitory to predominantly excitatory. This strongly implies that the intrinsic circuitry of the AC and/or how it is accessed by BA afferents undergo long-term plasticity [27, 29]. It also raises the possibility that pup odor may indeed be an important trigger for AC plasticity that has been commonly observed following maternal experience.

### Behavioral function of basal amygdala

The BA is activated during maternal behavior and in response to multimodal pup stimuli. Presenting mothers with either a hypothermic pup or the combination of pup odors and USVs results in significantly elevated *c-fos* expression in the BA relative to controls presented with no stimulus [14]. In that study, the synaptic targets of pup-responsive BA neurons were not identified, but the BA has been suggested to influence appetitive maternal behaviors (e.g. retrieval) through its projections to the NAcc and ventral pallidum [30, 31]. Our results here argue that, in addition to regulating reward circuitry, the BA might contribute to goal-directed maternal behaviors in a previously unappreciated manner by regulating the auditory processing of USVs. Interestingly, neurons in one of these other output pathways (BA-NAcc) showed no response to pup odors, raising the possibility of complexity in how different information regarding pup encounters may be private or shared with different BA targets.

Aside from its specific relationship to maternal behavior, recent evidence more broadly implicates the BA in regulating affiliative processes such as sociability and responses to social novelty [32, 33]. The basolateral amygdala (BLA) generally computes positive and negative valence signals [34] and emotional salience [35], and it is critical for motivated behavior. Distinct subsets of BLA neurons respond to either aversive or appetitive conditioned and unconditioned stimuli [36–39]. Each subset tends to preferentially access different downstream targets [40, 41]. Many BLA neurons exhibit positive (or negative) responses to multiple positive (or negative) stimuli, suggesting that they encode valence independent of the specific stimulus [42]. Less is known about activity in the basomedial amygdala, but it may be more closely linked to controlling behavioral aversion [43, 44]. Finally, analysis of ensembles embedded in a large population of simultaneously recorded BA neurons across diverse behavioral conditions show that these ensembles adopt distinct network configurations to encode long-term behavioral states such as social or spatial exploration [45, 46]

It is not known what computed quantities are represented by BA neurons that project to the AC, but all of the above observations reveal characteristics that they may share with other BA neurons. In any case, BA-AC neurons seem unlikely to carry straightforward sensory responses to odors. Given the known properties of the BA, they likely represent more abstract affective or state variables. We propose that the direct modulation of primary sensory activity with respect to these variables constitutes an underappreciated function of the amygdala.

### Future work

This work raises several interesting questions for future study. First, how are the population signals we detect here distributed among individual BA-AC neurons? Do they share the reported properties of other BA neurons? Recording from or imaging many identified individual neurons simultaneously during maternal behavior could help answer those questions. Second, are there state changes during maternal behavior such as arousal or engagement that could be in part modulated by the BA, affecting the responses to vocal signals? Future experiments might therefore focus on large-scale network dynamics in AC during maternal interactions and how they are altered by inputs from BA. Third, what aspects of maternal encounters are shared with other targets of BA, including reward circuits in the ventral pallidum and NAcc, and what effect do they have there?

## ACKNOWLEDGMENTS

The authors wish to thank J. Sturgill for technical assistance, the Zador, Tollkuhn and Li laboratories for their contributions, and Shea Lab members for helpful comments and discussion. Figures 1A-B, 3A-B, and 6A were created using Biorender.com. This work was supported by a grant to SDS from the National Institute of Mental Health (R01MH119250) and a grant to SDS from the C.M. Robertson Foundation

## AUTHOR CONTRIBUTIONS

Conceptualization: S.D.S. and A.C.N. Methodology: A.C.N., C.K., and S.D.S. Software: C.K. and S.D.S. Formal Analysis: A.C.N. and S.D.S. Investigation: A.C.N., J.C., and H.J.A. Writing – Original Draft: A.C.N. and S.D.S. Writing – Review and Editing: all authors. Visualization: A.C.N. and S.D.S. Supervision and Funding Acquisition: S.D.S.

## DECLARATION OF INTERESTS

The authors declare no competing interests.

## INCLUSION AND DIVERSITY

We support inclusive, diverse, and equitable conduct of research.

## METHODS

### Animals

All procedures were conducted in accordance with the National Institutes of Health’s Guide for the Care and Use of Laboratory Animals and approved by the Cold Spring Harbor Laboratory Institutional Animal Care and Use Committee. All experiments were performed on adult (aged 6-12 weeks) female mice that were maintained on a 12h:12h light-dark cycle and received food *ad libitum*. Throughout the text and figure legends, ‘n’ is used to refer the number of individual biological replicates. Due to the exploratory nature of our work, we did not a priori compute sample size for any of our experiments. Most mice were CBA/CaJ (Jax #000654), however neuroanatomical tracing experiments were performed in H2B-GFP *(Rosa26-stop^flox^-H2B-GFP,* gift from Bo Li), VGlut1Cre (*Slc17a7^tm1.1(cre)Hze^/J*; Jax #023527), VGlut2Cre (*Slc17a6^tm2(cre)Lowl^/J*; Jax #016963), or VGatCre (*Slc32a1^tm2(cre)Lowl^/J*; Jax #016962). We performed all behavioral experiments during the dark cycle.

### Retrieval behavior

Surrogates were generated by co-housing a virgin female with a pregnant CBA/CaJ dam beginning 1-5 days prior to birth. Pup retrieval behavior performance was first assayed on the day pups were born (P0) and again on P3 and P5 as described [6]. Briefly, retrieval behavior was performed in the home cage (39 cm by 20 cm by 16 cm), which was placed in a larger dark, sound-attenuated chamber (61 cm by 58 cm by 56 cm). The mother was removed and a surrogate was allowed to habituate to the behavior chamber in its home cage for 5 min with 5 pups in the nest. Pups were then removed for 2 minutes and subsequently returned to all four corners and the center. The trial started when the last pup was placed and each surrogate was given 5 minutes to retrieve all pups back to the nest. Videos were recorded in the dark under infrared light using a Logitech webcam (c920) with the IR filter removed.

Behavioral videos were annotated using BORIS (Behavior Observation Research Interactive Software) [47] by a trained observer who was blind to the experimental condition and day of testing. The observer manually scored the onset and offset of events including ‘search’, ‘retrieval success’, ‘retrieval error’, ‘investigation’, ‘air sniff’, and ‘nesting’, which were exported to MATLAB for further analysis. A normalized latency score was calculated using the following formula:

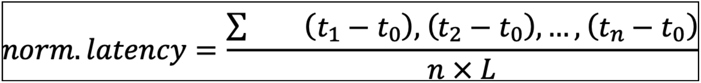

n = number of pups outside the nest, t0 = start of the trial, tn = time the nth pup was gathered, L = trial length (300s).

### MOE ablation

Following five days of maternal experience, surrogates were given an intraperitoneal (IP) injection of methimazole (MMZ; 50 mg/kg) (Millipore Sigma, #46429) dissolved in saline. We waited 7 days to allow sufficient time for the MOE to degenerate before retesting mice in the retrieval assay.

### Surgical procedures

Before all surgical procedures, mice were initially anesthetized with an IP injection (1.25 ml/kg) of an 80:20 mixture of ketamine (100 mg/ml) and xylazine (20 mg/ml) and were stabilized in a stereotaxic frame. Anesthesia was maintained throughout by vaporized isoflurane (1 – 2% as needed).

Depending on the specific experiment, we injected one or more adeno-associated viruses (AAVs) into the auditory cortex and/or the basal amygdala, as described in Results. See Table 1 for specific constructs used, titers, serotype, ordering information, and injected volumes. See Table 2 for injection coordinates. After the target volume of virus was expelled, the injection pipette was slowly retracted from the brain. In mice prepared for fiber photometry, a 5 mm length of 200 μm optical fiber (0.39 NA, Thorlabs) was implanted 200 μm above the BA injection site before being secured with Metabond dental cement along with a titanium head fixation bar. The scalp was sutured shut and the mouse was given a dose of meloxicam (2 mg/mL) for analgesia.

**Table 1.**

| *Viral construct* | *Ordering info* | *Volume and titer* |
| --- | --- | --- |
| AAVrg.CAG.tdTomato | Addgene<br>cat#: 59462-AAVrg | 0.45nL/injection site (6 total) of $1.3 \times 10^{13}$ GC mL<sup>-1</sup> |
| AAVrg.pmSyn1.EBFP.Cre | Addgene<br>cat#: 5107-AAVrg | 0.45nL of $5 \times 10^{12}$ GC mL<sup>-1</sup> |
| AAVrg.hSyn.Cre.WPRE.hGH | Addgene<br>cat#: 105553-AAVrg | 0.45nL of $1.2 \times 10^{13}$ GC mL<sup>-1</sup> |
| AAVrg.FLEX.tdTomato | Addgene<br>cat#: 28306-AAVrg | 0.45 nL of $7.5 \times 10^{12}$ GC mL<sup>-1</sup> |
| AAV5.syn.FLEX.GCaMP6s.WPRE | Addgene<br>cat#: 100845-AAV5 | 200nL of $2.9 \times 10^{13}$ GC mL<sup>-1</sup> |
| AAV9.synP.DIO.EGFP.WPRE.hGH | Addgene | 200nL of $4.3 \times 10^{13}$ GC |
|  | cat#: 100043-AAV9 | mL<sup>-1</sup> |
| AAV9.CAGGS.Flex.ChR2.tdTomato.WPRE.SV40 | UPenn | 200nL |

**Table 2.**

| *Brain Region* | *Coordinates* (measured from bregma unless noted) |
| --- | --- |
| Auditory Cortex | AP: -1.7, -2.1, -2.4mm<br>ML: 4.0mm<br>DV: -0.35, -0.55mm (from cortical surface)* |
| Basal Amygdala | AP: -1.5mm<br>ML: 3.0mm<br>DV: -5.4mm |

**Table 3.** Odorant Stimuli

| **Odorant** | **Sigma catalog #** |
| --- | --- |
| Mineral Oil | 330779 |
| Amyl Acetate | W504009 |
| Ethyl Tiglate | W246000 |
| R-Carvone | 124931 |
| S-Carvone | 435759 |
| Pup | N/A |
| Clean nest material | Bed r’Nest |

### Fiber photometry

Experiments were performed with a custom-built two-color fiber photometry system. The output from two LEDs (470nm/565nm; Doric) was focused into a fiber launch holding a 200 μm optical fiber (0.37 NA; Doric) that could be coupled to the implanted optical fiber via a lightweight, flexible cable during daily recording sessions. The LEDs were sinusoidally modulated 180 degrees out of phase at 211 Hz and emitted green and red light was collected from the optical fiber. Each color was separated and bandpass filtered and was detected by a dedicated photoreceiver (#2151, Newport Corporation). Light delivered to the brain was measured at ∼30 mW at the fiber tip.

Animals used in head-fixed recording sessions were habituated to fixation for 30-60 minutes once a day for 3-5 days prior to testing. At the start of each session, animals were connected to the patch cord and baseline signal was recorded for 2 minutes. Recordings from freely behaving mice were conducted identically with the exception that the cable attached to the head led to an optical swivel (Doric) to allow free movement. The signal from each photoreceiver was digitized at a sampling rate of 6100 Hz and acquired to a computer via a National Instruments DAQ device (NI-USB6211).

The raw data were used to calculate ΔF/F signals offline with custom written MATLAB code. First, the value at each peak in the sinusoidal signals was detected, resulting in an effective sampling rate of 211 Hz. Each signal was low pass filtered at 15 Hz and fit to a double exponential decay function, which was subtracted to correct for photobleaching during the trial. To correct for any movement artifact, we used a robust regression to compute a linear function for predicting the activity-independent component of the green signal based on the red. This component, which was in most cases small, was subtracted from the green signal. Finally, the result was mean subtracted and then divided by the mean to calculate ΔF/F. ΔF/F signals were Z-scored across all recording sessions from each mouse to compare calcium activity from multiple animals and days. The notation ‘Z-dFF’ is used here to distinguish Z-score normalized ΔF/F signals from raw ΔF/F signals (‘dFF’).

Four mice were excluded from the study as they did not exhibit fluctuations in their Ca^2+^ signal (likely due to misalignment in fiber placement). Individual trials were excluded if the Ca^2+^ trace contained unacceptably high contamination from artifact or if retrieval behavior was obscured. No subjects or trials were excluded for any other reason.

### Odor Presentation

Controlled presentation of odors to awake, head-fixed mice was achieved with a custom-built olfactometer as described [19]. Briefly, mice were head-fixed to a frame with their nose positioned in front of an odor port on a 10 cm diameter foam wheel. The wheel permitted the animal to freely walk or run or remain still. To deliver monomolecular odors, clean oxygen flow was briefly redirected by a solenoid assembly through the headspace of one of eight vials containing one odor (Table 5) diluted to 5% V/V in mineral oil. The odorized oxygen was mixed (1:10) with a clean air carrier stream to achieve a flow dilution of 0.5% saturated vapor. Pup and nesting material odors were presented by placing 3 pups or clean nesting material in a sealed chamber (4 oz mason jar) and swapping it with one of the odor vials in our olfactometer. Half of the pup chamber was kept on a warm pad and pup odor trials were limited to 10 minutes per session for the pups’ safety and comfort. Pups used for odor stimulation were used a maximum of 6 sessions per day and were returned to their mothers immediately following the experiment.

### USV recording and playback

Single pup vocalizations and tones were presented during electrophysiology recordings using one of the output channels of a National Instruments DAQ (NI-USB 6211) controlled by custom software written in MATLAB. Output was sent to an electrostatic speaker driver powering an electrostatic speaker (ED1/ES1, Tucker-Davis Technologies) positioned 4 inches behind the animal’s head. Stimuli were low pass filtered and amplified at 100kHz using a custom filter and preamp (Kiwa Electronics). A sound level meter (Model 407736, ‘A’ weighting; Ex-Tech) was used to calibrate the RMS for all stimuli to 70 dB SPL at the head. USVs were detected using a polarized condenser ultrasound microphone (CM16/CMPA, Avisoft Bioacoustics) placed 12” above the cage floor and were digitally sampled at 200 kHz via a National Instruments DAQ (NI-USB 6211) using custom software written in MATLAB.

### Electrophysiology

Prior to recording, the mouse was anesthetized with an 80:20 mixture (1.25ml/kg) of ketamine (100mg/ml) and xylazine (20 mg/ml). Extracellular recordings were made using the ‘loose patch’ method [5, 19]. Micropipettes were pulled from borosilicate glass filaments (O.D. 1.5mm, I.D. 0.86mm; BF150-86-10, Sutter Instrument) using a Flaming/Brown micropipette puller (P-97; Sutter Instruments) and back-filled with intracellular solution (125 mM potassium gluconate, 10mM potassium chloride, 2mM magnesium chloride, and 10mM HEPES pH 7.2) for a final resistance of 10 – 30 MΩ. Isolated single-unit neural activity was recorded using a BA-03X bridge amplifier (npi), low-pass filtered at 3 kHz, digitized at 10 kHz, and acquired using Spike2 software (v.7; Cambridge Electronic Design). Recording depth was measured by a piezoelectric micromanipulator (Sutter Instrument SOLO/E-116). All neurons were found from 300 μm – 1200 μm from the cortical surface, corresponding to approximately layers II – VI of auditory cortex.

Experiments were performed in an anechoic, sound-attenuated chamber (Industrial Acoustics). Single pup vocalizations and tones were presented to surrogates using one of the output channels on a Power1401 ADC/DAC board (Cambridge Electronic Design). Stimuli were low pass filtered at 100kHz and amplified using a custom filter and preamp (Kiwa Electronics). Output was sent to an electrostatic speaker driver powering an electrostatic speaker (ED1/ES1, Tucker-Davis Technologies) positioned 4 inches in front of the animal. All stimuli were calibrated to RMS of 65dB SPL at the animal’s head using a sound level meter (Ex-Tech, model 407736). Stimuli consisted of 7 log-spaced pure tones (16, 20, 25, 32, 40, 50, 64 kHz) or 8 natural pup calls recorded from CBA/CaJ mouse pups on postnatal day 2 (calls 1-4) or 4 (calls 5-8). Stimuli were presented for 100ms at an interstimulus interval of 4s.

### Optogenetic stimulation

To optogenetically activate BA terminals in the auditory cortex, the tip of an optical fiber (∅Ø 400 μm NA 0.39; Thor Labs) was positioned just over the craniotomy with a micromanipulator such that light shone directly on the cortical surface. Trains of light pulses were controlled by the timing output of an isolated pulse stimulator (A-M Systems) and consisted of 10 ms pulses of blue light (473nm; OEM laser) delivered at 20 Hz for 1 s [48, 49] and beginning 500 ms prior to the auditory stimulus. Laser power was 30 mW as measured at the tip of the optic fiber. Based on this, we estimate the power to be 2 – 10 mW/mm^2^ at a depth of 300-1000 mm from the cortical surface [50], where most recordings were made.

### Post-mortem histology and immunohistochemistry

At the end of each experiment, mice were delivered a lethal dose of pentobarbital (Euthasol), and then exsanguinated and transcardially perfused with PBS followed by 4% paraformaldehyde. In cases where the tissue was required for staining and/or microscopy, the brain and the main olfactory epithelium (MOE) were extracted and post-fixed overnight at 4°C. Brains were then transferred to a solution of 30% sucrose in PBS overnight at room temperature (RT) and subsequently sectioned on a freezing microtome at a thickness of 50µm. For visualization of GCaMP6s expression, brain sections were first incubated with a primary antibody raised against GFP in chicken (AVES) diluted 1:1000 in PDT (0.5% normal donkey serum, 0.1% Triton X-100 \in PBS) at 4 degrees overnight and then stained with a secondary anti-chicken Alexa 488 fluorophore raised in goat and diluted 1:500 in PDT for 2 h at room temperature. Nasal tissue was dissected from the skull [51] and sent to the CSHL histology core to be paraffin sectioned, H&E stained, and compared to a saline injected control (Figure 1). MOE ablation was confirmed by manual inspection of the processed tissue.

### Data analysis

All data analysis was performed with either Matlab (Mathworks) or Prism 9 software (GraphPad). Unless otherwise noted, data are reported as mean ± standard error.

Electrophysiology data were manually spike sorted into single unit spike trains with Spike2 (CED). All subsequent analyses were performed with custom-written code in MATLAB (Mathworks). Mean baseline firing rate was calculated as the mean spontaneous firing rate during a 1 s period just before each stimulus. This quantity was subtracted from the stimulus-evoked spike rate measured from 50 – 200 ms after the onset of each stimulus to calculate ‘response strength.’ Neurons that were presented fewer than 6 trials or had a mean firing rate of less than 0.5 spikes per trial were excluded from analysis, unless they exhibited clear auditory responses. To assess the statistical significance of responses to each auditory stimulus, we used a bootstrap procedure. For a response window length *t*, and a stimulus presented *n* times, we created a null distribution by computing the mean response strength across *n* windows of length *t*, randomly drawn from the full spike record and repeating this 10,000 times. Auditory-evoked response strengths in the top or bottom 2.5% of the null distribution were considered significantly excitatory or inhibitory auditory-evoked responses, respectively.

ROC analysis was used to assess whether the distribution of auditory-evoked response strengths from light-on trials were discriminable from light-off trials for a given neuron and stimulus. Analysis was performed in MATLAB (Mathworks) using a publicly available ROC curve function [52]. Because optogenetic modulation of auditory-evoked responses could be either enhancing or suppressive, we converted all auROC values < 0.5 to 1 minus their value.

To determine the degree to which optogenetic stimulation modified the firing response for each cell-stimulus pair that exhibited an auditory response, we used the following formula to calculate a modulation index. Here, RSaud equals the mean response strength to a given stimulus without optogenetic stimulation, and RSopto. equals the mean response strength in response to the same stimulus with optogenetic stimulation.

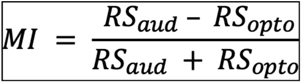

Optogenetic modulation of auditory responses was categorized as either enhancing or suppressive. This classification was based on whether the mean auditory-evoked response strength during optogenetic stimulation was higher or lower than the response strength evoked by the auditory stimulus alone.

For the analysis of odor-evoked activity, we calculated the mean amplitude of the z-dFF signal over the 4 s following the onset of odor presentation, and compared that to 2 s of baseline that occurred immediately before odor onset. We also calculated the difference between the mean signal at these two time points.

To examine whether an individual cell exhibited an evoked response to light stimulation alone (without auditory stimulus), we combined data from all light-only trials collected from a single cell. We then calculated the mean response strength within the 1s window of light stimulation and assessed whether the cell exhibited a statistically significant response based on the bootstrap procedure described earlier.

## Notes

### Competing Interest Statement

The authors have declared no competing interest.

### Summary of Updates

Significant changes: 1) There are now more data from additional animals strengthening the observation of olfactory responses in auditory cortically-projecting neurons in the basal amygdala to pups and arbitrary monomolecular odors. 2) In the previous version, we did not present odors to mice that were actively housed with pups out of concern for disrupting ongoing behavior assays. Responses to pups in experienced mice were therefore assessed in surrogates well past the 5 day course of co-housing. Thus, our data did not address whether responses to pup odor are maintained during the initial 5 days of surrogacy. We have now gathered those data and they reveal that robust responses to pup odor are maintained throughout this 5 day period. 3) This version includes an entirely new data set comparing the responses of auditory cortex projecting BA neurons (BA-AC) to pup odor with those of nucleus accumbens projecting BA neurons (BA-NAcc). We find that while BA-NAcc neurons respond to some common cage odors, only the BA-AC neurons respond to pup odor.

