## Supplementary material for "Multisensory integration of social signals by a pathway from the basal amygdala to the auditory cortex in maternal mice": Figures S1 - S6

### Supplementary Figure 1

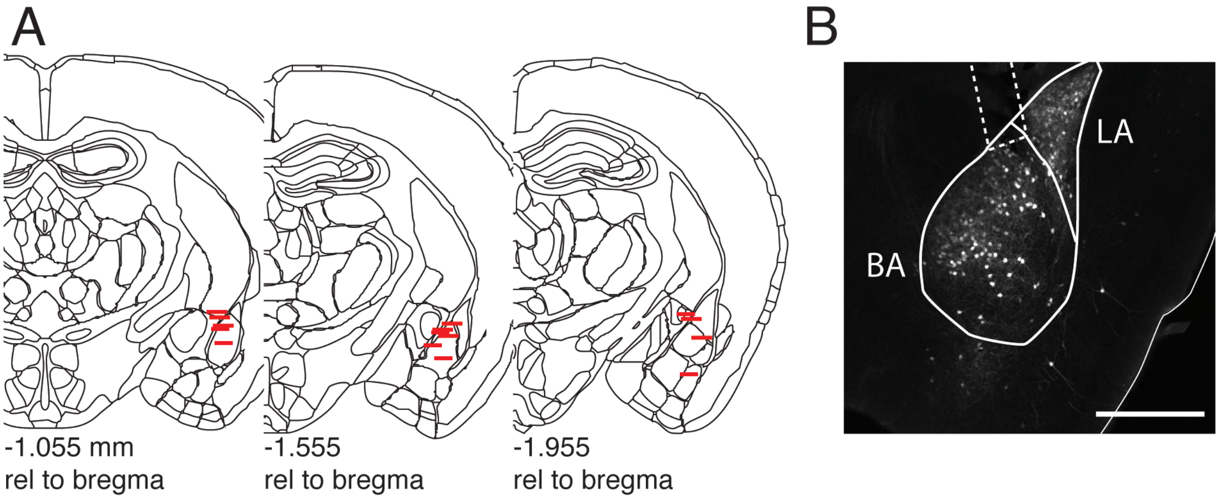


**Supplemental Figure 1:** Fiber implant map. (A) Schematic of coronal mouse brain sections depicting the location of optical fibers implanted in the BA (n = 15 mice; coordinates are measured from Bregma). (B) An example histological image in which the fiber track is depicted by the dashed line and the sections have been stained for GFP to visualize GCaMP6s expression within the BA (scale bar = 200 mM).

### Supplementary Figure 2

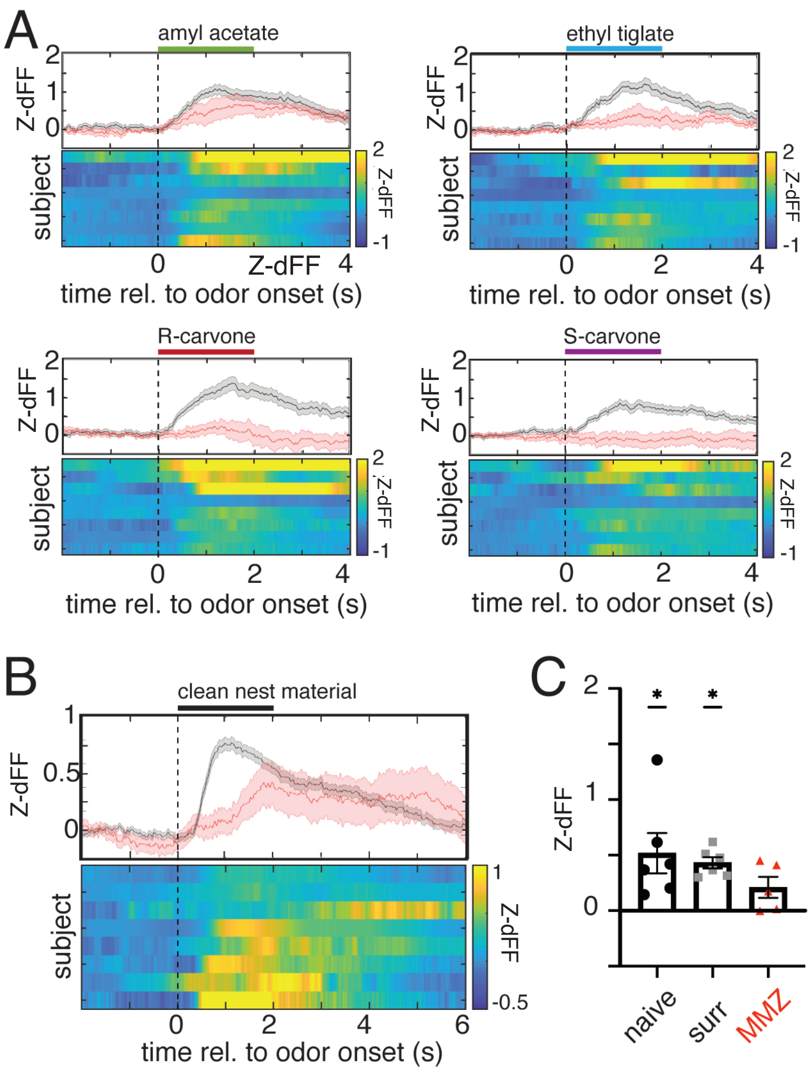


**Supplemental Figure 2:** Olfactory responses are variable across odors and mice. (A) Plots of the responses of all mice separately to each of four monomolecular odorants. In each plot, the upper panel is a mean trace of the responses of all mice to one monomolecular odor (gray) and the response to the same odor in a subset of those mice following ablation of the MOE with MMZ injection (red). The shaded area around each trace marks the SEM of the response. In the lower panel is a heatmap in which each row represents the mean response over multiple trials to one odor in one mouse, expressed as a Z score (Z-dFF), according to the color bar. Rows are ordered by the magnitude of each response. (B) Heatplot of mean responses to nest material for each mouse. Each row in the lower plot represents the mean response over multiple trials to nest material in one mouse, expressed as a Z score (Z-dFF), according to the color bar. Rows are ordered by the magnitude of each response. The upper plot shows a trace of the mean ± SEM response to nest material across all mice (gray) and the mean ± SEM response to nest material in the same mice following ablation of the MOE with MMZ injection. (C) Plot of mean response across all mice for nest material, comparing responses in naïve females, at the end of 5 d surrogacy, and after ablation of the MOE with MMZ treatment.  Each point denotes the mean response of one mouse, defined as the difference from pre-stimulus baseline. The distribution of responses to nest material was significantly different from 0 in naïve mice (n = 6 mice; 0.513 ± 0.181 z-dFF, Wilcoxon signed rank test, p = 0.031) and surrogates (n = 6 mice; 0.432 ± 0.051 z-dFF, Wilcoxon signed rank test, p = 0.031). Responses to nest material did not significantly differ from 0 after MMZ treatment (n = 5 mice; 0.210 ± 0.096 z-dFF, Wilcoxon signed rank test, p = 0.125).

### Supplementary Figure 3

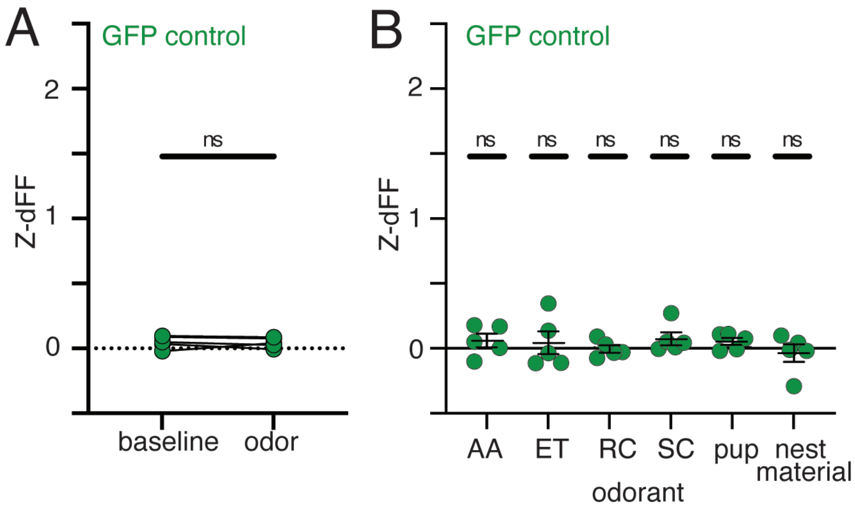


**Supplemental Figure 3:** Mice expressing activity-independent GFP in BA-AC show no detectable odor responses. (A) Plot of mean response across all mice for all odors, comparing baseline Z-dFF to mean Z-dFF during all odor presentations for mice that expressed GFP in BA-AC instead of GCaMP6s. Each pair of points indicates the mean fluorescence at baseline of one mouse and mean fluorescence during odor trials for the same mouse. These values were not significantly different (n = 5 mice; 0.04 ± 0.02 Z-dFF, paired t test, p = 0.815). (B) Plot of mean response across all GFP-expressing mice for four monomolecular odors (AA – amyl acetate, ET – ethyl tiglate, RC – R-carvone, SC – S-carvone), pup odor, and nest material. Each point denotes the mean response of one mouse, defined as the difference from pre-stimulus baseline (2 s). None of the distributions of responses for any of the odors were significantly different from 0 (n = 5 mice, Wilcoxon signed rank test, $p < 0.05$).

### Supplementary Figure 4

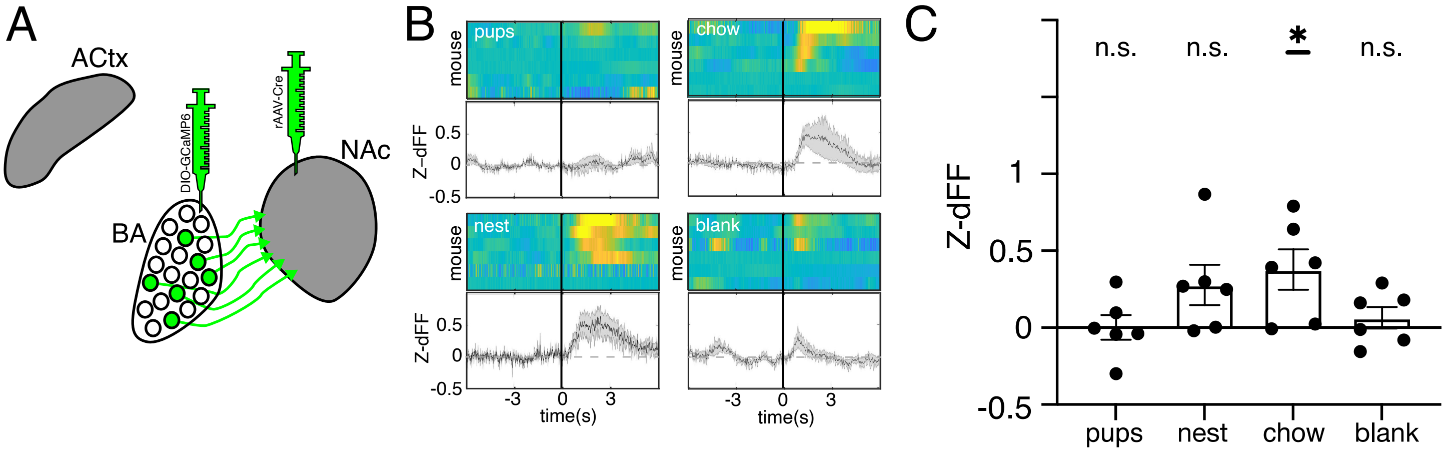


**Supplemental Figure 4:** BA neurons that project to nucleus accumbens show no response to volatile pup odors. (A) Schematic of our intersectional viral strategy for selectively labeling BA-NAcc projection neurons with GCaMP6s. (B) Plots of responses of all mice to volatile odors from pups, chow, clean bedding, and a blank stimulus. In each plot, the upper panel is a mean trace of the responses of all mice to the designated cage odor The shaded area around each trace marks the SEM of the response. In the lower panel is a heatmap in which each represents the mean response over multiple trials to one odor in one mouse, expressed as a Z score (Z-dFF), according to the color bar. Rows are ordered by the magnitude of each response. (C) Quantification of the increased Ca2+ signal in BA-NAcc during odor presentation. Generally, weak and inconsistent responses to odors were observed in most recorded subjects (n = 6 mice). All mice consistently exhibited no response to pups or the blank (empty vial) stimulus; several mice exhibited responses to bedding and chow, but when comparing mean responses of all mice to each odor with zero, only responses to chow achieved statistical significance: pups (0.002 ± 0.08), chow (0.38 ± 0.13), bedding (0.28 ± 0.13), and the blank (empty vial) (0.06 ± 0.07).

### Supplementary Figure 5

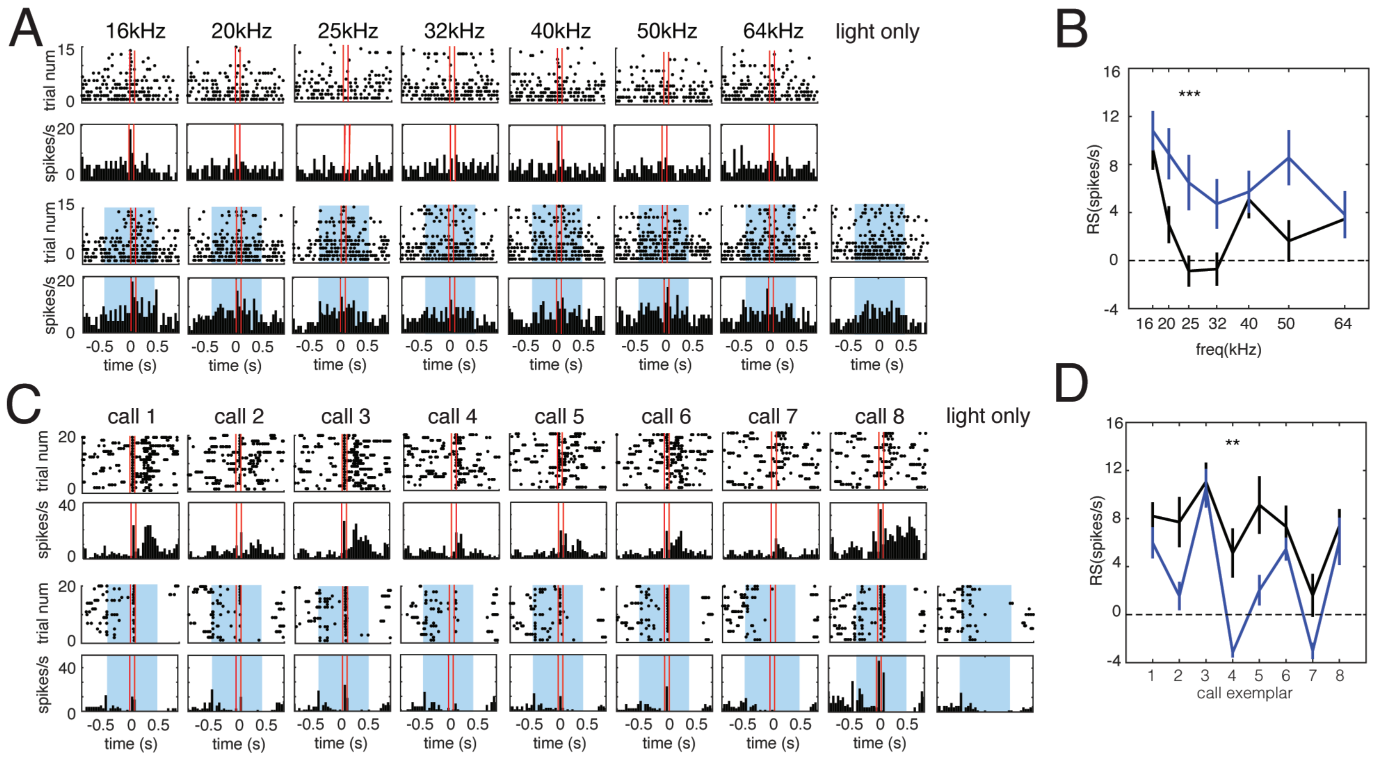


**Supplemental Figure 5:** Optogenetic activation of the BA-AC pathway elicits widespread and bidirectional modulation of auditory responses in the auditory cortex of surrogates. (A) Plots comparing responses of an auditory cortical neuron to logarithmically spaced pure tones when presented during light activation of BA-AC terminals and when presented alone. Data are from a surrogate female. Each stimulus is associated with a raster plot and peristimulus time histogram (bin size = 50 ms) from control trials (top row) and from light activation trials (bottom row with blue shading). The blue shading denotes the duration of the light train relative to the tone. Note that responses to the 25 kHz tone are significantly stronger when BA-AC terminals were activated by light (n = 15 trials; comparison of trials with and without light, unpaired t test with Bonferroni correction, $^{**}p < 0.01$). (B) Line plot of data from (A) comparing the mean baseline-subtracted firing rate evoked by each tone on control trials (black) and on light trials (blue). Vertical lines denote SEM. (C) Plots comparing responses of a different auditory cortical neuron to 8 different pup call exemplars when presented during light activation of BA-AC and when presented alone. Data are from a naïve female. Panels are organized as in (a). (D) Line plot of data from (C) comparing the mean baseline-subtracted firing rate evoked by each call on control trials (black) and on light trials (blue). Note that responses to call 4 are significantly weaker when BA-AC terminals were activated by light (n = 20 trials; comparison of trials with and without light, unpaired t test with Bonferroni correction, $^{***}p < 0.001$).

### Supplementary Figure 6

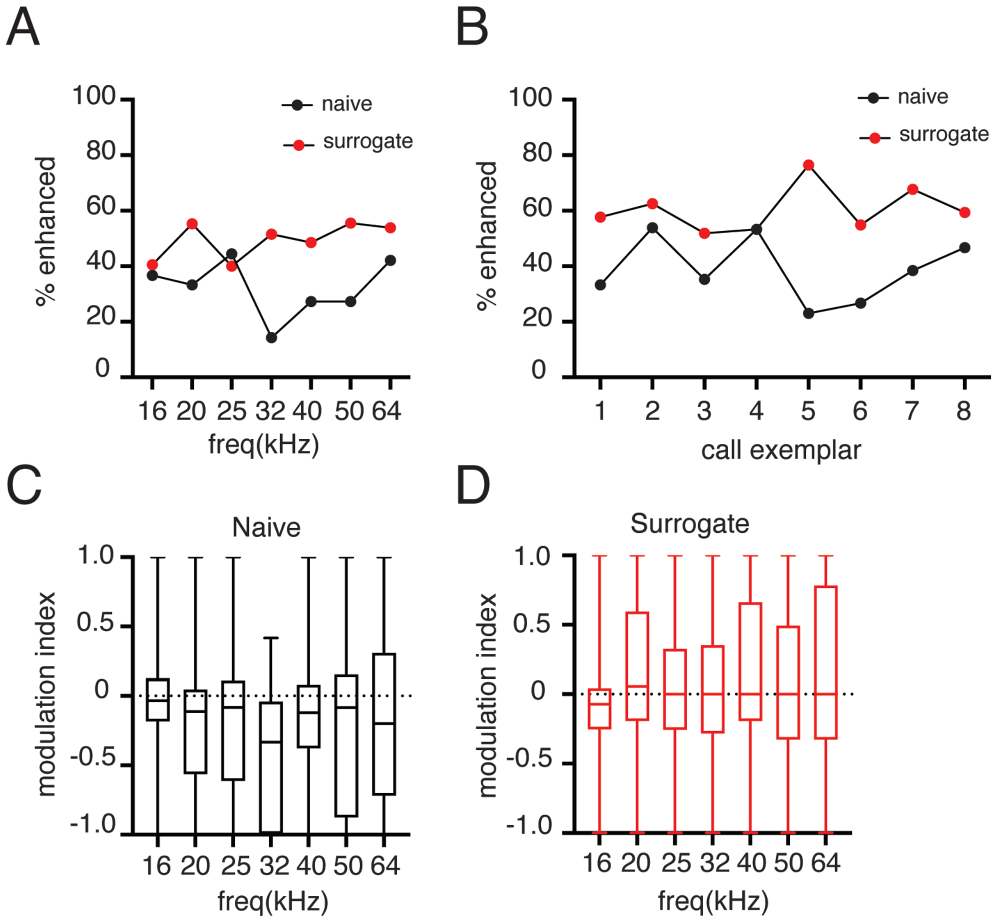


**Supplemental Figure 6:** Optogenetic modulation of auditory responses occurs independent of stimulus frequency. (A) The percentage of auditory responses enhanced by optogenetic activation is generally higher in surrogates compared to naïve animals. For the majority of stimulus frequencies this effect is not statistically significant when comparing the observed responses in ChR2 expressing animals to expected responses of GFP controls (Fisher's exact test, $p > 0.05$), with the exception of naïve responses to 32kHz frequency tones (Fisher's exact test, $p = 0.020$). (B) The same is true for call exemplars with the exception of surrogate responses to call exemplar 8 (Fisher's exact test, $p = 0.0003$). (C) The average modulation index across units (n = 31-33) recorded from naïve mice (n = 7) is not significantly different across frequencies (2 way ANOVA, stimulus source of variation, $p = 0.0569$). (D) The same is true of units (n = 36-44) recorded from surrogates (n = 10, 2 way ANOVA, stimulus source of variation, $p = 0.6677$).
